# The RNA-dependent interactions of phosphatidylinositol 4,5-bisphosphate with intrinsically disordered proteins contribute to nuclear compartmentalization

**DOI:** 10.1101/2024.03.19.585734

**Authors:** Martin Sztacho, Jakub Červenka, Barbora Šalovská, Ludovica Antiga, Peter Hoboth, Pavel Hozák

**Author notes:** Corresponding authors: Pavel Hozák, Martin Sztacho. These authors contributed equally to this work (Martin Sztacho, Jakub Červenka, Barbora Šalovská).

## Abstract

The RNA content is crucial for the formation of nuclear compartments, such as nuclear speckles and nucleoli. Phosphatidylinositol 4,5-bisphosphate (PIP2) is found in nuclear speckles, nucleoli and nuclear lipid islets and is involved in RNA polymerase I/II transcription. Intriguingly, the nuclear localization of PIP2 was also shown to be RNA-dependent. We therefore investigated whether PIP2 and RNA cooperate in the establishment of nuclear architecture. In this study, we unveiled the RNA-dependent PIP2-associated (RDPA) nuclear proteome in human cells by mass spectrometry. We found that intrinsically disordered regions (IDRs) with polybasic PIP2-binding K/R motifs are prevalent features of RDPA proteins. Moreover, these IDRs of RDPA proteins exhibit enrichment for phosphorylation, acetylation and ubiquitination sites. Our findings reveal that RDPA protein BRD4 associates with PIP2 in an RNA-dependent manner via electrostatic interactions, and that elevated PIP2 levels increase the number of BRD4 protein nuclear foci. Thus, we propose that PIP2 spatiotemporally orchestrates nuclear processes through association with RNA and RDPA proteins and affects their ability to phase separate. This suggests pivotal role of PIP2 for the establishment of a functional nuclear architecture competent for gene expression.

## Introduction

Differentially phosphorylated inositol headgroups of different phosphoinositides (PIPs) serve as a recognition code for the recruitment of a plethora of interacting proteins (1–4). PIPs are typically embedded in eukaryotic cell membranes where they regulate processes such as vesicular trafficking, actin polymerization or autophagy (5,6). The most abundant nuclear PIP is phosphatidylinositol 4,5-bisphosphate (PIP2). PIP2 localizes to nuclear speckles, nucleoli and small nucleoplasmic structures called nuclear lipid islets (NLIs) (7,8). Nuclear PIPs are involved in gene expression (2,7,9). In particular, nuclear PIP2 regulates transcription by affecting the condensation capacity of the RNA Pol2 initiation complex (10). Interestingly, nuclear compartments containing PIP2, such as the aforementioned nuclear speckles, nucleoli and nucleoplasmic transcription initiation foci, are formed by the process of phase separation (11–17).

The phase separation-driven formation of membraneless compartments, sometimes referred to as ‘biomolecular condensates’, is associated with enhanced kinetics of biochemical reactions in the living cell (18–21). The formation of these compartments provides high local concentrations of reaction components and forms diffusion barriers that serve as adsorption catalyst surfaces (22). In addition, biomolecular condensates allow the sequential progression of processes through the successive coupling of subsequent reactions in multilayered compartmentalized reaction chambers, such as ribosomal biogenesis in nucleoli, packaging of hnRNP particles in Cajal bodies, or the involvement of nuclear speckles in pre-mRNA splicing (13,23–25).

The formation of phase-separated biomolecular condensates can be mathematically described and computationally modelled using the theory of stickers and linkers (26). Stickers are local modules that allow multivalent intra- and intermolecular interactions and are represented by classical globular domains of proteins or by stretches of charged amino acids connected by flexible linker regions in intrinsically disordered regions (IDRs) (27,28). Condensation of IDR-containing proteins is often driven by charged amino acid stretches within unstructured protein regions (29–31). In addition, it has previously been described that changes in net charge and amino acid types within an IDR can even navigate proteins to different core regions (29). Thus, both the amino acid composition and the posttranslational modifications (PTMs), which together generate the charge pattern of the IDR, are important determinants of sub-nuclear protein localization (31). Electrostatic interactions appear to be fundamental determinants of condensate properties (32,33). Thus, negatively charged polymeric molecules such as RNA are the important factors in the formation and dissolution of some nuclear condensates.

Indeed, the biomolecular condensates in the nucleus are typically formed by low-affinity multivalent interactions between proteins and RNA (34,35). RNA has a positive or negative effect on condensate formation, depending on the situation and the type of RNA (36–38). The short RNA molecules buffer and thus reduce the local tendency to form a condensate. Conversely, longer RNA molecules often increase condensate formation (36,37,39,40). In addition, PTMs such as phosphorylation are another important regulatory step affecting condensate formation or dissolution (14,29,36,41). The interaction between higher-order folded RNA and lipid molecules has been suggested previously (17,42–48). Higher-order RNA has a scaffolding function that brings together RNA-binding proteins to form nuclear subcompartments (49–53). The formation of these RNA folds depends on intra- and intermolecular double-stranded RNA (dsRNA) duplexes. However, a general mechanism or identification of common mechanistic principles has been lacking.

We hypothesized that negatively charged nuclear PIPs are interesting candidates for the regulation of biomolecular condensation via phase separation. PIPs provide a platform for the recruitment of interacting proteins, thereby increasing their local concentration. RNA and PIPs may cooperate in the formation of condensates, such as in a process of influenza virus particle biogenesis (54). PIPs carry a negative charge, which could ultimately alter the overall net charge of condensates and thus influence condensate formation and size. Therefore, the possible spatial interplay between nuclear PIPs and RNA in regulating condensation seems plausible. Indeed, a recent study showed that not only proteins and RNA, but also metabolites including phospholipids (e.g. PIPs) are enriched in condensates *in vitro* (55). We have previously shown that RNA is important for nuclear PIP2 levels, as RNA removal by RNase A dramatically decreased the PIP2 signal measured by immunofluorescence (7). In the current study, we speculate that the higher-order RNA might be responsible for the correct localization of PIP2 in the eukaryotic nucleus. Therefore, we used bacterial RNase III, normally associated with siRNA processing, to remove short dsRNA regions followed by quantitative mass spectrometry (MS) proteomic analysis of the nuclear fraction. We identified the RNA-dependent PIP2-associated (RDPA) nuclear proteome, followed by bioinformatic analyses of the physicochemical properties of the identified proteins. Subsequently, we experimentally tested our model in which nuclear PIP2 serves as a recognition motif, regulates the phase separation capacities of interacting proteins, and thus plays a major role in the establishment of nuclear architecture.

Our results revealed the molecular mechanism by which PIP2 acts as a molecular wedge for the recruitment of lysin/arginine (K/R) motif-containing RDPA proteins by structured RNA molecules, leading to local regulation of condensation potential. The level of nuclear PIP2 therefore dictates where particular RDPA proteins accumulate and potentially form condensate. Thus, changes in RDPA proteins localization and condensation potential might affect the rates of the biochemical reactions involved. Taken together, our data explain formation of the specific type of biomolecular condensates via interactions between RNA, proteins and PIP2, and are therefore relevant to our understanding of the principles underlying the establishment of functional nuclear compartments.

## Material and Methods

### Cell culture, antibodies

HeLa (ATCC no. CCL2) cells were cultured in suspension in DMEM media (Sigma D6429) supplemented with 10% fetal bovine serum in spinner flasks at 37 °C 10% CO_2_ atmosphere for 5 days. U2OS (ATCC no. HTB96) were grown in DMEM media (Sigma D6429) with 10% FBS at 37 °C in a humidified 5% CO_2_ atmosphere. Antibodies were used in this study at concentrations according to the manufacturer’s instructions (Table 1).

**Table 1:** Antibodies used in this study.

| Antibody | Vendor | Ref number | Immunofluorescence | Western Blot |
| --- | --- | --- | --- | --- |
| Anti-PIP2 | Echelon Biosciences Inc., USA | AB010220-28 | 5 µg/mL |  |
| Anti-BRD4 | Sigma-Aldrich, St. Louis, MO, USA | HPA015055 | 0.20 µg/mL | 0.20 µg/mL |
| Anti-SON | Sigma-Aldrich, St. Louis, MO, USA | HPA062997 | 1 µg/mL |  |
| Anti-PIP5KA | Sigma-Aldrich, St. Louis, MO, USA | HPA029366 |  | 0.10 µg/mL |
| Anti-SHIP2 | Abcam, UK | Ab70267 |  | 0.50 µg/mL |
| Anti-actin | Sigma-Aldrich, St. Louis, MO, USA | MABT219 |  | 1.7 µg/mL |
| Anti-GST | Abcam, UK | Ab6613 |  | 2 µg/mL |
| IRDye 800 CW Donkey anti-Rabbit IgG | LI-COR Biosciences, Lincoln, NE, USA | 926-32213 |  | 0.10 µg/mL |
| IRDye 800 CW Donkey anti-Mouse IgG | LI-COR Biosciences, Lincoln, NE, USA | 926-32212 |  | 0.10 µg/mL |
| IRDye 680RD Donkey anti-Mouse IgG | LI-COR Biosciences, | 926-68072 |  | 0.10 µg/mL |
|  | Lincoln, NE,<br>USA |  |  |  |
| IRDye 680RD Donkey anti-Rabbit IgG | LI-COR<br>Biosciences,<br>NE, USA | 926-68073 |  | 0.10 µg/mL |
| Goat anti-Mouse IgG (H+L)<br>Highly Cross-Adsorbed, Alexa<br>Fluor 568 | Invitrogen, MA,<br>USA | A-11031 | 5 µg/mL |  |
| Goat anti-Rabbit IgG (H+L)<br>Highly Cross-Adsorbed, Alexa<br>Fluor 488 | Invitrogen, MA,<br>USA | A-11034 | 5 µg/mL |  |

### Immunofluorescence labelling

U2OS cells were grown on high-performance cover glasses of 12 mm in diameter with restricted thickness-related tolerance (depth = 0.17 mm ± 0.005 mm) and the refractive index = 1.5255 ± 0.0015 (Marienfeld 0107222). The cells were fixed with 4% formaldehyde for 20 min, permeabilized with 0.1% Triton X-100 for 5 min and three times washed in phosphate-buffered saline (PBS). Specimens were blocked by 5% bovine serum albumin (BSA) in PBS for 30 min. The specimens were incubated 1 h with primary antibodies (Table 1), three times washed with PBS and subsequently incubated 30 min with respective secondary antibodies (Table 1). Followed by three PBS washes and one wash with tap water, the specimens were air-dried for 20 min and mounted in 90% glycerol with 4% n-propyl gallate (NPG) media.

### Nuclear RNA extraction

Suspension HeLa cells were grown in 500 mL spinner flasks in DMEM media and harvested by 800 g centrifugation for 15 min at 4 °C. HeLa cell pellets were resuspended in 3 mL of lysis buffer (50 mM Hepes pH 7.4, 150 mM NaCl, 2 mM MgCl_2_, 0.5% NP-40) with 20 U RNase inhibitor (Applied Biosystem, MA, USA, S17857). Sample was spun down at 1000 g, 4 °C for 5 min. Supernatant representing cytoplasmic fraction was taken out. Additional 3 mL of lysis buffer were added and spun at 1000 g, 4 °C for 1 min and supernatant was discarded. Additional 1.2 mL of lysis buffer was added to pellet and mixed. After that step, 2.5 mL of TRIzol (Sigma, MO, USA, BCCF2003) and 2.5 mL of chloroform were added into the sample and vortexed vigorously. Sample was centrifuged at 12,000 g for 5 min at 4 °C. The upper phase was taken out and transferred to a new tube, 0.7 volume of isopropanol was added and mixed. The sample was centrifuged at 12,000 g for 15 min at 4 °C. Supernatant was removed, the pellet was washed by 80% ethanol and aspirated. The pellet was air dried for 10 min, dissolved in water and RNA concentration was measured by NanoDrop. The DNA was cleaved out by incubation with DNase I enzyme, when 160 µg of RNA was mixed with 12 µL of supplied reaction buffer and 30 U of DNase I (Sigma, MO, USA, D4527). The reaction was incubated for 30 min at 37 °C, then mixed with 0.8 volume of isopropanol and centrifuged at 12,000 g for 15 min at 4 °C. Pellet was washed with 80% of ethanol, air-dried for 10 min at RT, and dissolved in RNA-free water. The concentration of RNA was determined by NanoDrop measurement.

### RNase III treatment of semi-permeabilized U2OS cells

To obtain semi-permeabilized cells, 90% confluent U2OS cell cultures were washed twice with PBS. Cells were permeabilized by 0.1% Triton X-100 at RT in buffer (20 mM Hepes pH 7.4, 150 mM NaCl, 25% glycerol, 1 mM DTT, cOmplete EDTA-free protease inhibitor cocktail (La Roche Ltd., Basel, Switzerland, 05056489001)), and treated for 10 min at RT by RNase III enzyme (2 U of RNase III (Short Cut, New England BioLabs, Massachusetts, USA, M0245S) with 20 mM MnCl_2_ in 1× reaction ShortCut buffer). This step was followed by five times wash in PBS and fixation by 4% PFA for 15 min at RT. Cells were than permeabilized by 0.2% Triton X-100 for additional 15 min at RT. After three PBS washes, the cells were blocked for 30 min in 3% BSA in PBS and subjected to immunofluorescence protocol. Measured integral signal density of nuclear PIP2 signal was analyzed by FIJI software (56). Data were obtained from three biological replicates, in total 76 cells were quantified for non-treated control cells condition, and 89 cells were quantified for RNase III treated cells condition.

### Colocalization evaluation of immunofluorescence

Colocalization of epitopes was evaluated by the image analysis carried out using the Coloc2 plugin in FIJI software calculating three different correlation coefficients as suggested in (57). The nuclear outlines were segmented manually. Degree of colocalization was determined by the Pearson’s correlation coefficient, Spearman’s rank correlation value, and Manders’ correlation coefficients (M1 and M2) of the signals from the two analyzed channels. The significance of each statistical analysis was determined by the Student’s t-tests. The randomized images were obtained as described in (57).

### Nuclear lysate preparation

One liter of suspension culture of HeLa cells was spun at 1300 g at 4 °C for 15 min. Pellet was resuspended in 7 mL of buffer (50 mM Hepes pH 7.4, 150 mM NaCl, 1 mM DTT, cOmplete (La Roche Ltd., Basel, Switzerland, 05056489001)) and subjected to 20 strokes in Dounce homogenizer. Cell nuclei were sedimented by 1800 g centrifugation at 4 °C for 5 min. Supernatant was collected as cytoplasmic fraction. Nuclear pellet was washed four times in 10 mL of buffer. Clean nuclear pellet was sonicated in Soniprep 150 (MSE, London, UK) bench top sonicator (1 s on, 1 s off for 30 cycles at power of 10 microns amplitude). Sonicated lysate was spun down at 13,000 g at 4 °C for 15 min. Supernatant was collected as nuclear fraction. Protein concentration was determined by Pierce BCA Protein Assay (Thermo Fisher Scientific, Waltham, MA, USA, 23227) according to the manufacturer’s protocol.

### Preparation of PLCδ1 PH domain

For expression and purification of GST-tagged recombinant proteins PLCδ1 PH domain (1-140 aa) wild type and R40A mutation of PLCδ1 PH domain in pGST5 were used plasmids constructed previously (58). The PIP-binding protein domains were expressed in 100 mL of BL21 (DE3)-pLysS *E. coli* (Stratagene, Santa Clara, USA) culture. Transformed cells were incubated for approximately 4 h at 37 °C until OD = 0.6. Expression was then induced by 0.1 mM IPTG for additional 2 h. Samples were lysed by sonication with Soniprep 150 (MSE, London, UK) benchtop sonicator (4 s on, 4 s off for 1 min at power 10 microns of amplitude) in ice-cold buffer (50 mM Hepes pH 7.5, 150 mM NaCl, 1 mM DTT, cOmplete (La Roche Ltd., Basel, Switzerland, 05056489001)) and spun down at 13,000 g at 4 °C for 15 min. Supernatants were used for purification of recombinant proteins by 2 h incubation with GST-agarose beads at 4 °C according to manufacturer’s protocol (Sigma Aldrich, St. Louis, USA, G4510). SDS-PAGE electrophoresis was used to check level of expression and purity of purification.

### Pull-down assay with PIP2-conjugated beads at various treatment conditions

Four mL of nuclear lysate from suspension culture of HeLa cells at protein concentration of 2.5 mg/mL were prepared in buffer (50 mM HEPES, pH 7.4, 150 mM NaCl, 2 mM MgCl_2_, 1 mM DTT, cOmplete (La Roche Ltd., Basel, Switzerland, 05056489001) and PhosStop RNase inhibitors (La Roche Ltd., Basel, Switzerland, 4906837001)). Two types of agarose beads were used in pull-down assays: control beads, P-B000 (Echelon Biosciences Inc., UT, USA) and PI(4,5)P2-conjugated beads, P-B045A (Echelon Biosciences Inc., UT, USA). Forty µL of beads slurry were added into 650 µL of nuclear lysate per condition and incubated overnight at 4 °C. To test the biochemical nature of PIP2-BRD4 interaction, the following treatments were used in the respective specimens as indicated in Figure 4A: the addition of 30 µg of nuclear RNA extract, 300 mM NaCl, 100 mM NH_4_OAc, 10% 1,6-hexanediol, and 10% dextran. The beads were washed three times with 1 mL of ice-cold buffer and spun down at 800 g at 4 °C for 5 min. The supernatant was discarded, and the beads were boiled in 30 µL of Laemmli buffer for 10 min. The beads were spun down, and the supernatant was loaded into the SDS-PAGE gel. After trans-blotting, the membranes were blocked with 3% BSA for 30 min. The membranes were washed with 0.5% Tween 20/PBS for 15 min. The dilution of the primary antibody (Table 1) was prepared in 3% BSA/PBS and incubated for 2 h. Secondary antibody was used according to manufacturer’s instruction (Table 1). Western blot (WB) signals at each pull-down condition in every repetition were normalized to the highest signal (the PIP2 beads with RNA added condition). Statistical analysis was performed using Student’s t-tests on four replicates.

### Isolation of natural PIP2-structures and testing of BRD4-association

Four mL of nuclear lysate from suspension culture of HeLa cells at protein concentration 2.5 mg/mL were prepared in buffer (50 mM HEPES, pH 7.4, 150 mM NaCl, 2 mM MgCl_2_, 1 mM DTT, cOmplete (La Roche Ltd., Basel, Switzerland, 05056489001) and PhosStop RNase inhibitors (La Roche Ltd., Basel, Switzerland, 4906837001)). Two types of GST agarose beads were used in the pull-down assays: GST-PLCδ1 PH domain (1-140 amino acids, wild type) and R40A mutation of GST-PLCδ1 PH domain. Twenty µL of beads slurry were added into 650 µL of nuclear lysate per condition and incubated overnight at 4 °C. To test biochemical nature of BRD4 association with PIP2-containing structures, the following treatments were used in respective specimens as indicated in Figure 4B: the addition of 30 µg of nuclear RNA extract, 300 mM NaCl, 100 mM NH_4_OAc, 10% 1,6-hexanediol, and 10% dextran. The beads were washed three times with 1 mL of ice-cold buffer and spun down at 800 g at 4 °C for 5 min. The supernatant was discarded, and the beads were boiled in 30 µL of Laemmli buffer for 10 min. The beads were spun down, and the supernatant was loaded into the SDS-PAGE gel. After trans-blotting, the membranes were blocked with 3% BSA for 30 min. The membranes were washed by 0.5% Tween 20/PBS for 15 min. The dilution of the BRD4 primary antibody was prepared in 3% BSA/PBS and incubated for 2 h. Secondary antibody was used according to manufacturer’s instruction (Table 1). WB signals at each pull-down condition in every repetition were normalized to the highest signal of the wild type GST-PLCδ1 PH domain with RNA added condition. Statistical analysis was performed using Student’s t-tests on six replicates.

### PIP2-conjugated beads pull-down assay with spiked recombinant GST-PLCδ1 PH domain

Two mL of nuclear lysate from suspension culture of HeLa cells at protein concentration of 2.5 mg/mL were prepared in buffer (50 mM HEPES, pH 7.4, 150 mM NaCl, 2 mM MgCl_2_, 1 mM DTT, cOmplete (La Roche Ltd., Basel, Switzerland,05056489001) and PhosStop RNase inhibitors (La Roche Ltd., Basel, Switzerland, 4906837001)). Two types of agarose beads were used in pull-down assays: control beads, P-B000 (Echelon Biosciences Inc., UT, USA) and PI(4,5)P2-conjugated beads, P-B045A (Echelon Biosciences Inc., UT, USA). Forty µL of beads slurry were added into 650 µL of nuclear lysate per condition and incubated overnight at 4 °C. To test specificity of the effect of RNA on PIP2 binding of BRD4 protein the addition of 30 µg of nuclear RNA extract and 1 µg of purified recombinant soluble GST-PLCδ1 PH domain (1-140 amino acids, wild type), were used in the respective specimens as indicated in Figure 4C. The beads were washed three times with 1 mL of ice-cold buffer and spun down at 800 g at 4 °C for 5 min. The supernatant was discarded, and the beads were boiled in 30 µL of Laemmli buffer for 10 min. The beads were spun down, and the supernatant was loaded into the SDS-PAGE gel. After trans-blotting, the membranes were blocked with 3% BSA for 30 min. The membranes were washed with 0.5% Tween 20/PBS for 15 min. The dilutions of the primary antibodies (anti-GST, anti-BRD4) were prepared in 3% BSA/PBS and incubated for 2 h. Secondary antibodies were used according to manufacturer’s instruction (Table 1). WB signals at each pull-down condition in every repetition were normalized to the highest signal of the PIP2-beads upon RNA extract addition condition. Statistical analysis was performed using Student’s t-tests on four replicates.

### Mass spectrometry experimental pipeline

One mL of nuclear fraction from suspension culture of HeLa cells with protein concentration of 2.5 mg/mL was used per condition in three independent biological replicates. Twenty-five µL of washed wild-type or R40A mutation GST-PLCδ1 PH domain immobilized to glutathione agarose beads were added into the respective reactions and incubated at 4 °C for 1.5 h while rotating to allow PIP2–PH domain interaction. The RNase III-treated samples were pre-incubated with 4 U of RNase III (New England Biolabs, Ipswich, Massachusetts, USA, M0245S) in reaction supplemented by 20 mM MnCl_2_ at 15 °C for one hour. These samples were incubated with wild-type GST-PLCδ1 PH domain. After the incubation of all samples, the beads were centrifuged at 300 g at 4 °C for 2 min. Supernatant from each sample was carefully discarded. Beads were spun down and washed twice in 1 mL of buffer (50 mM Hepes, pH 7.4, 150 mM NaCl, 1 mM DTT, cOmplete (La Roche Ltd., Basel, Switzerland, 05056489001)) and subjected to sample preparation for MS measurement. Beads were resuspended in 100 mM triethylammonium bicarbonate (TEAB) containing 2% sodium deoxycholate (SDC). Proteins were eluted and cysteines were reduced in one step by heating with 10 mM final concentration of Tris-(2-carboxyethyl)phosphine (TCEP; 60 °C for 30 min). Beads were removed by centrifugation, and proteins in the supernatant were incubated with 10 mM final concentration of methyl methanethiosulfonate (MMTS; 10 min RT) to modify reduced cysteine residues. In-solution digestion was performed with 1 µg of trypsin at 37 °C overnight. After digestion, the samples were centrifuged, and the supernatants were collected and acidified with trifluoroacetic acid (TFA, final concentration of 1%). SDC was removed by ethylacetate extraction (59). Peptides were desalted using homemade stage tips packed with C18 disks (Empore) according to Rappsilber et al. (60).

One liter of suspension culture of HeLa cells was spun at 1300 g at 4 °C for 15 min. Pellet was resuspended in 7 mL of buffer (50 mM Hepes pH 7.4, 150 mM NaCl, 1 mM DTT, cOmplete (La Roche Ltd., Basel, Switzerland, 05056489001)) and subjected to 20 strokes in Dounce homogenizer. Cell nuclei were sedimented by 1800 g centrifugation at 4 °C for 5 min. Supernatant was collected as cytoplasmic fraction. Nuclear pellet was washed four times in 10 mL of buffer. Clean nuclear pellet was sonicated in Soniprep 150 (MSE, London, UK) bench top sonicator (1 s on, 1 s off for 30 cycles at power of 10 microns amplitude). Sonicated lysate was spun down at 13,000 g at 4 °C for 15 min. Supernatant was collected as nuclear fraction. Three independent biological replicates per sample were prepared for MS analysis as described above.

### Liquid chromatography-tandem mass spectrometry (LC-MS/MS) analysis

Nano reversed phase column (EASY Spray column, 50 cm, 75 µm ID, PepMap C18, 2 µm particles, 100 Å pore size) was used for LC-MS/MS analysis. The mobile phase A was composed of water and 0.1% formic acid. The mobile phase B was composed of acetonitrile and 0.1% formic acid. The samples were loaded onto the trap column (Acclaim PepMap 300, C18, 5 µm, 300 Å, 300 µm, 5 mm) at 15 µL/min for 4 min. The loading buffer was composed of water, 2% acetonitrile, and 0.1% TFA. Peptides were eluted with the mobile phase B gradient from 4% to 35% in 60 min. Eluting peptide cations were converted to gas phase ions by electrospray ionization and analyzed on a Thermo Orbitrap Fusion (Q OT qIT, Thermo Fisher Scientific, Waltham, MA, USA). Survey scans of peptide precursors from 350 to 1400 m/z were performed at 120 K resolution (at 200 m/z) with a 5×10^5^ ion count target. Tandem MS was performed by isolation at 1.5 Th with the quadrupole, HCD fragmentation with normalized collision energy of 30, and rapid scan MS analysis in the ion trap. The MS/MS ion count target was set to 10^4^, and the max injection time was 35 ms. Only those precursors with charge state 2–6 were sampled for MS/MS. The dynamic exclusion duration was set to 45 s with a 10 ppm tolerance around the selected precursor and its isotopes. Monoisotopic precursor selection was turned on. The instrument was run in top speed mode with 2 s cycles (61).

### Raw data processing

Raw data files acquired by LC-MS/MS were processed with MaxQuant v1.6.11.0 (62). Peak lists were searched against the human SwissProt database (May 2020) using Andromeda search engine (63). Minimum peptide length was set to seven amino acids, and two missed cleavages were allowed. Dithiomethylation of cysteine was set as a fixed modification, while oxidation of methionine and protein N-terminal acetylation were used as variable modifications. Only peptides and proteins with false discovery rate (FDR) lower than 0.01 were accepted. Protein intensities were normalized using MaxLFQ algorithm (64). MaxQuant output data were further analyzed using Perseus v1.6.13.0 (65) and visualized in R v4.0.0 (66). Briefly, protein groups identified at the 0.01 FDR level were further filtered to remove potential contaminants, decoys, and proteins identified based on modified peptides only. The resulting matrix was filtered based on the number of missing values (100% of valid values in at least one of the groups), and after log2 transformation, missing values were imputed from normal distribution (width = 0.3 times standard deviation (SD) and shift = 1.8 times SD of the original distribution). One replicate from the samples enriched using the wild-type GST-PLCδ1 PH domain-conjugated beads was identified as an outlier with an overall lower MS intensity and removed from the downstream analysis.

### Identification of nuclear RNA-dependent PIP2-associated (RDPA) proteins

A two-step statistical analysis was performed to identify the RNA-dependent PIP2-associated (RDPA) proteins. In the first step, the “PIP2-associated proteome” was identified by comparing the proteome enriched using the wild-type GST-PLCδ1 PH domain-conjugated beads to the proteome enriched using the point mutation (R40A) GST-PLCδ1 PH domain-conjugated beads; a one-sided Student’s t-test was performed with a permutation-based FDR correction. In the second step, the significant proteins (FDR < 0.05, S0 = 0.2, n = 195 protein groups) were then compared to the samples enriched using the wild-type PLCδ1 PH domain-conjugated beads from samples pretreated with RNase III. A two-sided Student’s t-test with a permutation-based FDR correction was applied to identify differentially enriched (FDR < 0.05, S0 = 0.2) protein groups after RNase III treatment (n = 183; downregulated n = 168). In both comparisons, Student’s t-test was performed in Perseus v1.6.13.0 (Supplementary Table 1). The results were visualized using the R package “ggplot2”.

### Manipulation of PIP2 level in cells

MISSION esiRNA (Sigma-Aldrich, USA, EHU114801-20UG) was used to deplete human PIP5K1A. MISSION esiRNA (Sigma-Aldrich, USA, EHU081051-20UG) was used for depletion of the human SHIP2. MISSION siRNA Universal Negative Control #1 (Merck, NJ, USA, SIC001) was used as the negative control. U2OS cells were seeded 24 h before the transfection on 12 mm in diameter glass coverslips with restricted thickness-related tolerance (depth = 0.17 mm ± 0.005 mm) and the refractive index = 1.5255 ± 0.0015 (Marienfeld, 0107222) at 70% confluency. The cells were transfected using Lipofectamine RNAiMax (Invitrogen, MA, USA, 13778150) for 24 h according to manufacturer’s protocol and subjected to immunofluorescence staining. Protein depletion efficiency was confirmed by WB assay (Supplementary Figure 16) and quantification of changes in nuclear PIP2 levels was performed based on the IF signal. The images were acquired as Z stacks at Leica STELLARIS 8 FALCON (Leica Mikrosysteme Vertrieb GmbH, Wetzlar, Germany) confocal microscope with × 63 oil objective NA 1.4.

### Image analysis of BRD4 protein foci

Identification of foci was performed using a macro pipeline written in FIJI software (56). Briefly, the channels from the Z-stack acquisition were split and analyzed in sequence. First, the nuclear area detected by the DAPI signal was identified. The nuclear area was further processed using the 3D Gaussian blur function. Next, the channel with visualized protein was processed by Gaussian blur 3D on the ROI previously identified as the nuclear area and the outside was deleted. The 3D object counter was then used to identify the protein foci using a minimum size filter of 10 voxels. The results showed the number of foci per cell and their volume in μm^3^. Statistical analysis was performed using the Kolmogorov-Smirnov test to analyze the frequency distributions of the raw volumetric data.

### Functional characterization of the RDPA proteome using Metascape

The functional analysis of the RDPA proteome (proteins associated with PIP2 in RNA-dependent positive manner) was performed by the Metascape tool using the default settings (67). The protein list for Metascape analysis comprised of 168 proteins; one majority protein ID was selected per protein group (Supplementary Table 17). The analysis was performed against a default Metascape background set. The enriched functional terms were identified using a default Metascape algorithm using a hypergeometric test. The significant terms were then hierarchically clustered into a tree based on Kappa-statistical similarities among their gene memberships. Then 0.3 kappa score was applied as the threshold to cast the tree into term clusters. We then selected a subset of representative terms from this cluster and converted them into a network layout. More specifically, each term is represented by a circle node, where its size is proportional to the number of input genes belonging into the term, and its color represents its cluster identity (i.e., nodes of the same color belong to the same cluster). Terms with a similarity score > 0.3 are linked by an edge (the thickness of the edge represents the similarity score). The network was visualized with Cytoscape (v3.1.2) with “force-directed” layout. One term from each cluster was selected to have its term description shown as a label. The complete results from the Metascape analysis are provided in Supplementary Table 18.

### Data preparation for bioinformatic analyses

Majority protein IDs from significant protein groups (see Raw data processing) were mapped to the UniProtKB database (68) (*Homo sapiens*, Swiss-Prot, reference proteome UP000005640, release 2022_01) and their canonical protein sequences were obtained. Seven datasets were used for further analyses: i) proteins associated with PIP2 in higher order RNA-dependent positive manner (called RDPA proteins, n = 183); ii) proteins quantifiable in at least two replicates of nuclear fraction, but not in cytosolic fraction, with RDPA proteins added (called Nucleo-specific proteins, n = 848); iii) proteins quantifiable in at least two replicates of nuclear fraction, with RDPA proteins added (called Nuclear fraction proteins, n = 3,655); iv) proteins quantifiable in at least two replicates of cytosolic fraction, but not in nuclear fraction (called Cytosol-specific proteins, n = 428); v) proteins quantifiable in at least two replicates of cytosolic fraction (called Cytosolic fraction proteins, n = 3,379); vi) all proteins quantifiable in at least two replicates of cytosolic or nuclear fraction, with RDPA proteins added (called Total cell proteome, n = 4,082) and vii) reference human proteome from UniProtKB with one protein sequence per gene (called Reference proteome, n = 20,577). Complete list of employed protein IDs and the graphical illustration of overlaps between original datasets are in Supplementary Table 3, 4 and Supplementary Figure 1.

### Search for protein domains binding to PIP2

Protein domains with ability to bind to PIP2 were selected based on previously published data (69–72). The PROSITE database (73) and InterPro database (74) were used to search for proteins with such features in Swiss-Prot database (*Homo sapiens*, reference proteome UP000005640, release 2022_02), resulting in 1,552 distinct proteins. This protein list was then compared with our datasets (Supplementary Table 5).

### RNA-binding capability

Protein IDs from datasets were compared to the list of all human proteins with experimental evidence for RNA-binding, according to the RNAct database (75) (3,717 reviewed proteins from Swiss-Prot, mapped to the reference proteome UP000005640) (Supplementary Table 6, 8).

### Association with phase separation

Protein IDs from datasets were compared to the list of all human proteins connected with phase separation or membraneless organelles, according to PhaSepDB2.0 (76) (4,014 reviewed proteins from Swiss-Prot, mapped to the reference proteome UP000005640) (Supplementary Table 7, 8).

### Prediction of intrinsically disordered regions (IDRs)

Disordered regions were predicted by ESpritz (version 1.3) (77) with three prediction types: X-Ray, Disprot, and NMR and decision threshold 5% False Positive Rate (Supplementary Table 9). For further analysis, R-based script (R, version 4.3.1) (66) was created, which uses UniProtKB accession numbers and search them against the Database of Disordered Protein Predictions (D2P2) (78). However, not all UniProtKB accession numbers were successfully mapped to D2P2. On average, we were able to retrieve the information for 95.6% (from 94.5% to 96.5%) of the input protein sequences for most of the datasets. The exception was Reference proteome, where we were able to retrieve information for 88.9% of the input protein sequences. Protein was counted as IDR-containing protein only, if at least one IDR with minimum length of 20 amino acid residues was predicted in this protein. The pI and hydrophobicity (GRAVY) of the disordered regions were calculated using the R package “peptides”, functions pI and hydrophobicity.

### K/R-rich motifs abundance analysis

Short sequence motifs rich for lysine and/or arginine (K/R motifs) were analyzed as described previously (79). Briefly, K/R motifs: [KR]-x(3,7)-K-x-[KR]-[KR], [KR]-x(3,7)-K-x-[KR] and [KR]-x(3,7)-K-x-K were searched in all datasets using ScanProsite tool (80), match mode set as greedy, no overlaps (Supplementary Table 10).

### Analysis of K/R motifs enrichment in IDRs

The analysis was performed using a custom R-based script (R, version 4.1.3), which uses information about presence of K/R motif in dataset from ScanProsite tool, and search UniProtKB accession numbers of such proteins against D2P2. K/R motif was counted as present in IDR only, if at least three predictors from the D2P2 predicted IDR with minimum length of 20 amino acid residues at the site of the K/R motif (Supplementary Table 11).

### Analysis of function of K/R motifs in IDRs

Localization of K/R motifs (i.e., [KR].{3,7}K.[KR][KR], [KR].{3,7}K.[KR] and [KR].{3,7}K.K) in proteins to structured regions or IDRs and gene ontology (GO) analyses of proteins containing such motifs were assessed with SLiMSearch4 tool (81), with disorder score cut-off set to 0.95 (Supplementary Table 19-27). The results of the analysis were visualized using “bubble plots”. In the plots, the y-axis shows the –log10 adjusted p-value of proteins from a GO category, the x-axis shows the log2 enrichment factor. The size of the bubble corresponds to the number of proteins.

### PTM proximity analysis

A database of known posttranslational modification (PTM) sites was downloaded (August 6, 2022) from the PhosphoSitePlus database (82). For the following known PTMs: acetylation, methylation, phosphorylation, sumoylation, and ubiquitination, we explored whether they are located in the IDR containing the K/R motifs identified by abovementioned analysis of K/R motifs enrichment in IDRs using a custom R-based script (Supplementary Table 16).

### Statistical analyses and data visualization

Statistical relevance of depletion or enrichment of particular feature (e.g., IDR content) between datasets was analyzed in R, version 4.3.1 (66) using hypergeometric test (function phyper). All plots were generated using the R package “ggplot2”. In the boxplots, the bold line indicates the median value; box borders represent the 25th and 75th percentiles, and the whiskers represent the minimum and maximum value within 1.5 times of interquartile range. Outliers out of this range are depicted using solid dots.

### Data availability

The mass spectrometry data and result tables have been deposited to the ProteomeXchange Consortium via the PRIDE (83) partner repository with the dataset identifier PXD045745. The R code used in this study is available at GitHub (https://github.com/BarboraSal/RNA_dependent_PIP2_associated_nuclear_proteome).

## Results and discussion

### RNA is important for PIP2 nuclear localization

RNA is the critical integral element for the coherence of many membraneless structures (84,85). Indeed, RNA is important in regulating the phase separation of proteins forming condensate assemblies (86). The scaffolding RNA typically adopts higher order folds that are enabled by the formation of dsRNA regions (87,88). We therefore hypothesized that dsRNA structures are important for PIP2 localization in the nucleus. To test this hypothesis, we used RNase III treatment of semi-permeabilized cells followed by immunofluorescence labelling of PIP2 and the nuclear speckles marker protein SON. RNase III does not have a conserved target sequence but recognizes dsRNA structures (89,90). The nucleus is a very dense environment and nuclear PIP2 forms sub-diffraction limited foci, so we used super-resolution microscopy (91–93). We observed that RNase III-mediated RNA cleavage greatly reduced the total nuclear PIP2 signal, whereas based on SON staining, the structural integrity of nuclear speckles is not completely abolished (Figure 1A-C). These results suggest that the nuclear localization of PIP2 is dependent on the presence of higher order dsRNA in specific regions. Thus, this observation postulates an intimate relationship between PIP2 and RNA in the nucleus.

**Figure 1:**
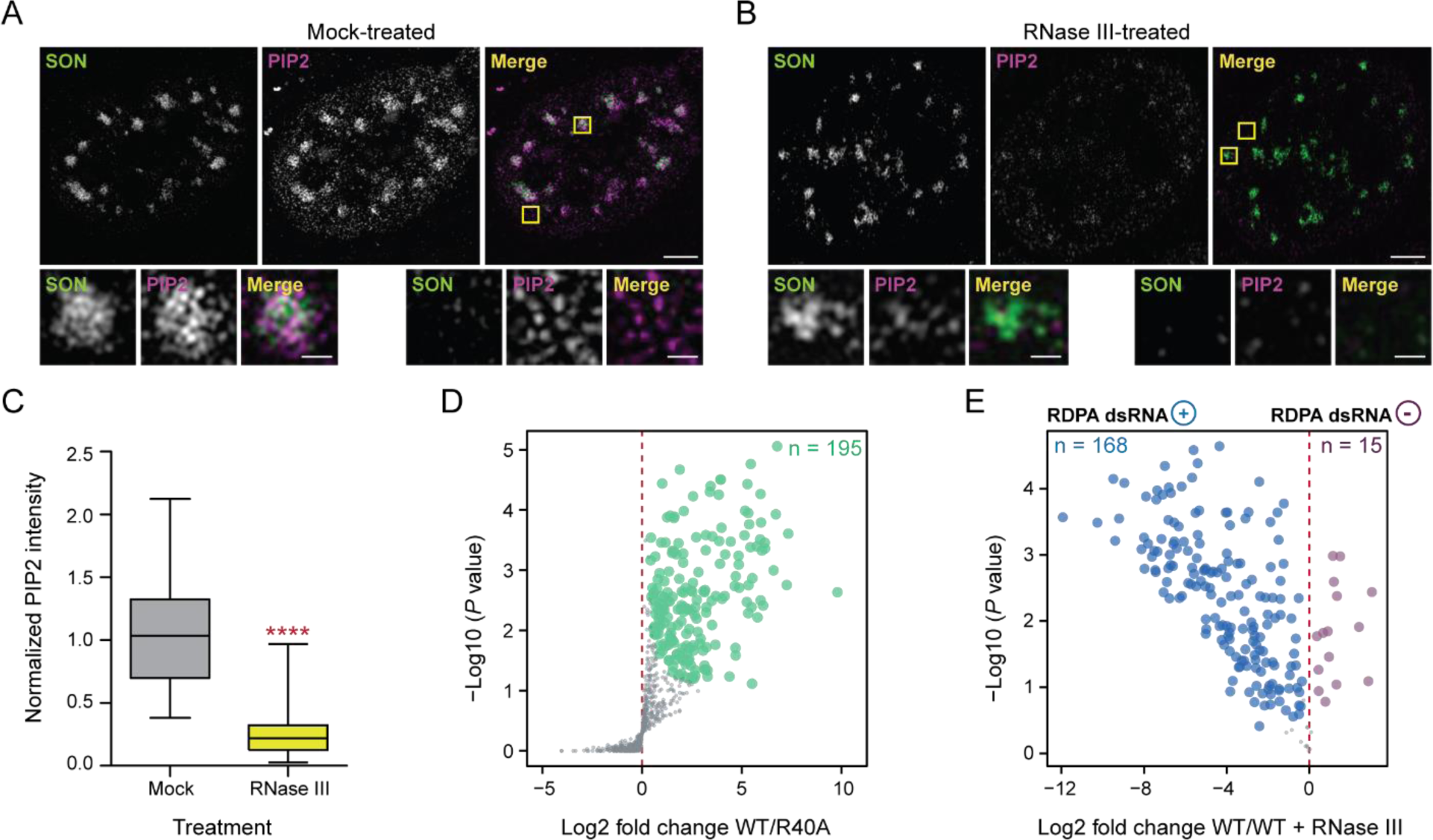
Identification of the RNA-dependent PIP2-associated (RDPA) nuclear proteome. **(A-B)** The effect of RNase III treatment on PIP2 and SON localization in cell nuclei visualized by immunofluorescence staining and super-resolution microscopy. U2OS cells were semi-permeabilized, treated by RNase III, and subsequently stained with PIP2 and SON-specific antibodies. Images were acquired by structured illumination microscopy (SIM). **A)** Mock-treated U2OS cell nucleus, **B)** RNase III-treated nucleus with detailed zoom-in insets of PIP2 and SON localization in the nucleoplasmic and nuclear speckle subcompartments. Scale bars correspond to 5 µm and 1 µm in the image and inset details, respectively. **C)** The quantification of normalized mean PIP2 signal intensity. Statistical analysis was performed using Student’s t-tests (**** P < 0.0001), n = 3, N = 76 cells mock-treated, N = 89 RNase III-treated cells. **D)** Volcano plot shows the PIP2-associated nuclear proteome identified by MS analysis; the significantly enriched protein groups (n = 195) are depicted in green (FDR < 0.05 & S0 = 0.2). **E)** Volcano plot of proteins whose PIP2-structures association is regulated by the presence of dsRNA. Protein groups showing a statistically significant (FDR < 0.05 & S0 = 0.2) loss (n = 168; RDPA dsRNA +, positively regulated by dsRNA) or gain (n = 15; RDPA dsRNA -, negatively regulated by dsRNA) of association with PIP2 after dsRNA cleavage are shown in blue or purple, respectively. A Student’s t-test with a permutation-based FDR correction was performed using a function provided in the Perseus software.

### Identification of RNA-dependent PIP2-associated nuclear proteome

We searched for proteins that are important for the formation of nuclear architecture dependent on the interplay between RNA and PIP2. Considering the importance of RNA in the formation of nuclear subcompartments, it has been suggested to use RNases to identify proteins involved in the formation of nuclear structures produced by phase separation (88). We aimed to identify the proteins that associate with PIP2-containing structures in an RNA-dependent manner. Therefore, we designed a label-free quantitative MS approach based on phospholipase C pleckstrin homology (GST-PLCδ1 PH) domain pull-downs from nuclear lysates treated or untreated with RNase III. The wild-type GST-PLCδ1 PH domain binds PIP2 with high specificity, whereas its point mutation R40A abolishes this binding capacity (58). In the first step of this experimental workflow, we prepared nuclear lysates suitable for comparative MS as described previously (79). The wild-type and R40A GST-PLCδ1 PH domain variants were then attached to agarose beads via a glutathione S-transferase (GST) tag and incubated with the nuclear lysates. Analysis of MS data of proteins bound to both domains with differential abundance led to the identification of the PIP2-associated nuclear proteome (Figure 1D). In parallel, we digested dsRNA in the third nuclear lysate sample with RNase III prior to the wild-type GST-PLCδ1 PH domain pull-down followed by MS measurement. This step resulted in the depletion of 168 protein groups and the enrichment of 15 protein groups, allowing us to identify PIP2-associated proteins that were differentially changed upon dsRNA cleavage (Figure 1E). Thus, this experimental pipeline allowed us to identify the nuclear RNA-dependent PIP2-associated (RDPA) proteome. Our data show that the vast majority of PIP2-associated proteins lost their PIP2 association upon dsRNA removal.

PIP2 is normally embedded in cytoplasmic membranes where it regulates various processes through interactions with a variety of proteins with PIPs binding domains. We hypothesized that nuclear PIP2 might also be involved in the recognition and binding of various proteins, thereby regulating the localization of their actions. Such proteins typically contain canonical PIPs binding domains with well-defined globular structures, e.g. pleckstrin homology (PH), phox homology (PX), Fab-1, YGL023, Vps27 and EEA1 (FYVE) domains, etc. In order to assess the abundance of PIPs binding domains in the RDPA proteome, we generated in parallel several reference datasets by analyzing the nuclear and cytosolic fractions isolated from the same HeLa cell line using the same LC-MS/MS proteomic pipeline (see M&M section for more details). We then searched for these domains in the RDPA proteome and in datasets of proteins identified exclusively in the nucleus (’nucleo-specific’, 848 proteins) and in the cytosol (’cytosol-specific’, 428 proteins). In addition, we generated the ‘total cell proteome’ (4,082 proteins) by combining all proteins identified in our MS analyses of nuclear and/or cytosolic proteomes. For comparison, we also provide the analysis of the additional proteomes, i.e. ‘nuclear fraction’ proteins, ‘cytosolic fraction’ proteins and ‘reference proteome’ (Supplementary Tables 3, 4 and Supplementary Figure 1; see M&M for details). However, in agreement with previously published data (2,79), the frequency of canonical PIPs binding domains was very low in both RDPA sub-populations. Only 11 out of 183 RDPA proteins positively regulated by the presence of dsRNA contain the PIPs binding domains (Supplementary Table 5), suggesting that these domains are not a major route of PIP2 association. Interestingly, none of the proteins negatively regulated by the presence of dsRNA possess such a domain. Notably, this dataset (16 proteins) that showed increased PIP2 binding upon dsRNA cleavage is rather small for reasonable statistical correlation with other datasets. Therefore, we focused further analyses only on RDPA proteins with a positive dsRNA effect on their PIP2 binding, and unless otherwise noted, the term RDPA proteins refers to this subpopulation.

### RDPA proteome is enriched for proteins with phase separation capacity and PIP2-binding motifs in their IDRs

Nuclear PIP2 localizes to two archetypal liquid-like structures formed by phase separation - nuclear speckles and nucleoli (13,21,25). The appropriate subcompartmentalized structure of such multiphase arrays depends on the presence of architectural RNAs (23,51,87,94,95). Furthermore, there is a higher concentration of RNA in the nucleus compared to the cytosol, suggesting that RNA is the critical driving force for the preferential formation of such structures in the nucleus. Our bioinformatic analyses confirmed the enrichment of RNA-binding properties within the RDPA proteome (Figure 2A & Supplementary Figure 2A, Supplementary Table 6), a predictable feature due to the RNase III treatment step in our MS workflow. Importantly, we showed that the RDPA proteome is the most enriched for proteins with the ability to phase separate from all datasets (Figure 2A; Supplementary Figure 2A, Supplementary Table 7). This search was based on the overlap of proteins from the RDPA proteome with the PhaSEP database, which contains proteins associated with phase separation and membraneless organelles (96). Finally, the RDPA proteome is enriched for proteins with both RNA binding and phase separation capabilities together (Figure 2A; Supplementary Figure 2A, Supplementary Table 8), suggesting that these two properties are linked, as described elsewhere (39).

**Figure 2:**
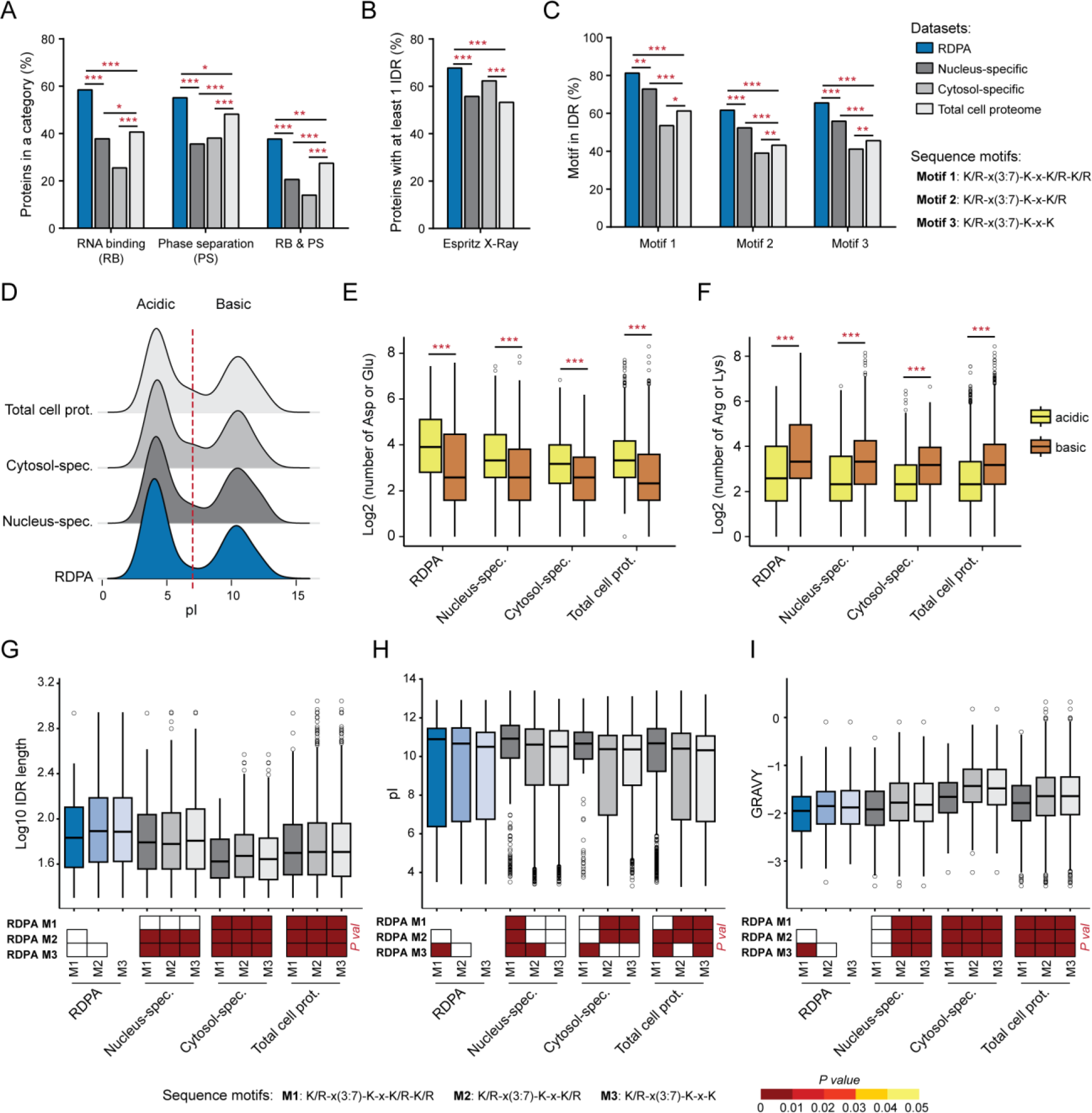
Bioinformatic analyses of RDPA proteome features. **A)** RDPA proteome is significantly enriched for RNA binding, phase separation capacity, and combination of both properties. **B)** RDPA proteome is enriched for IDRs longer than 30 amino acid residues predicted by ESpritz X-Ray. **C)** Percentage of PIP2-binding K/R motif sites localized in IDRs (from all K/R motif sites in the dataset) is elevated in RDPA proteome (only IDRs predicted by at least three different predictors with minimal length 20 amino acid residues were considered). Statistical analysis was performed using a hypergeometric test (* P < 0.05, ** P < 0.01, and *** P < 0.001). **D)** pI values of IDRs predicted by ESpritz X-Ray in the analyzed protein datasets show a bimodal distribution. **E)** The IDRs in the “acidic” population (pI < 7) are enriched with D/E amino acid residues. **F)** The IDRs in the “basic” population (pI > 7) are K/R-rich. Statistical analysis was performed using a Wilcox test (*** P < 0.001). **G-I)** IDRs containing the three K/R motifs tend to be significantly longer **(G)**, more basic **(H)**, and more hydrophilic **(I)** in the RDPA proteome compared to other datasets. Statistical analysis was performed using a pairwise Wilcox test with Benjamini-Hochberg correction. Datasets: RDPA – proteins with positive dsRNA effect on their PIP2 binding (dsRNA+), Nucleo-specific – proteins identified exclusively in the nucleus, Cytosol-specific – proteins identified exclusively in the cytosol, and Total cell proteome – all proteins identified in the nucleus and/or cytosol.

Phase separation is often mediated by multivalent interactions between proteins and RNA (35,38,86). One of the typical structural features with phase separation capacity are IDRs. It has been shown that the nuclear proteome is enriched for proteins containing IDRs (97–99), suggesting that nuclear proteins are prone to phase separation and thus to the formation of biomolecular condensates. Indeed, our bioinformatic analysis revealed that the RDPA proteome is significantly enriched for proteins containing IDRs among other datasets (Figure 2B, Supplementary Table 9). These results were confirmed using three different IDR predictors, which yielded similar results (Supplementary Figure 2B, C).

Previously described K/R motifs have been identified as regions important for PIP2 interaction (2,79). We therefore screened our datasets for the presence of three known K/R motifs - K/R-x(3,7)-K-x-K/R-K/R, K/R-x(3,7)-K-x-K/R and K/R-x(3,7)-K-x-K. These motifs were abundant in the RDPA proteome, but only the K/R-x(3,7)-K-x-K/R-K/R motif (the longest one) was significantly enriched compared to all other datasets (Supplementary Figure 3A,B, Supplementary Table 10). Since the RDPA proteome is enriched for K/R motifs and RNA-binding proteins, and its PIP2 association is RNA-dependent, it can be assumed that not all RDPA proteins interact directly with PIP2. It is therefore possible that positively charged K/R motifs serve as binding sites for negatively charged RNA molecules. Different K/R motif lengths suggest localization to different nuclear loci and thus involvement in different processes. This effect is likely to be manifested by different K/R content providing different affinity and thus retention time in a particular nuclear compartment as shown elsewhere (29).

We then investigated whether these K/R motifs are localized in proteins inside or outside the predicted IDRs. In the case of the RDPA proteome, all three motifs had a significantly increased abundance in IDRs (Figure 2C; Supplementary Figure 3C, Supplementary Table 11). Interestingly, the longest, and thus least permissive motif to search, showed the highest frequency in IDRs (four times higher than outside IDRs). Based on the above data, the RDPA proteins contain PIP2-binding K/R motifs within their IDRs and possess phase separation and RNA binding capacity.

### RDPA proteins contain long hydrophilic IDRs with acidic D/E-rich and basic K/R-rich regions

The aforementioned K/R motifs within IDRs are typical examples of multivalent interaction modules employed in the phase separation-driven formation of biomolecular condensates (29–31). To determine the distribution of net charge in the IDRs, we analyzed the isoelectric points (pI) of all predicted IDRs in the RDPA proteome. We used the Database of Disordered Protein Predictions (78), and all nine different IDR predictors showed a similar bimodal distribution pattern of IDRs (only IDRs with a minimum length of 20 amino acid residues were considered) based on their pI (Figure 2D; Supplementary Figure 4, 5A). The acidic peak (pI < 7) represents IDRs enriched in D/E amino acid residues (Figure 2E; Supplementary Figure 5B, 6A, C). The second peak represents basic IDRs (pI > 7) enriched for K/R amino acid residues (Figure 2F; Supplementary Figure 5C, 6B, D). These data show that IDRs have a bimodal pI distribution regardless of their cell fraction origin. Interestingly, the D/E motif has been described as important for the negative regulation of protein condensation capacity (100,101). Thus, PIP2-mediated recruitment of RDPA IDRs containing the D/E-rich regions could have a negative effect on condensation and limit the size of condensates formed. Therefore, PIP2 may function in defining the local concentration of nuclear RDPA proteins and regulating their localization and condensation capacity as suggested in (55).

We demonstrated that K/R motifs are more abundant within predicted IDRs than in external structured regions (Figure 2C). Therefore, we focused our further analysis on IDRs (predicted by at least three different predictors with a minimum length of 20 amino acid residues) containing the K/R motifs. In particular, we evaluated the average length of K/R motif-containing IDRs across different datasets. The results show that RDPA proteins have significantly longer IDRs than other datasets, regardless of the K/R motif type analyzed (Figure 2G, Supplementary Figure 7, 8, 9, Supplementary Table 12). Importantly, the RDPA and nucleo-specific proteomes are specifically significantly enriched for two IDR types with average lengths of ∼300 and ∼800 amino acid residues (Supplementary Figure 9B, Supplementary Table 13). Longer IDRs are more prone to condensation due to an increased degree of intrinsic disorder (102), i.e. a higher number of disordered amino acid residues (Supplementary Figure 10, Supplementary Table 9).

Analyzing the pI distribution of K/R motif-containing IDRs confirmed the bimodal distribution of pI found in IDRs (Figure 2D), irrespective of the K/R motif present (Supplementary Figure 11B). However, the pI of K/R motif-containing IDRs is more basic than acidic (Supplementary Figure 11A), consistent with the higher abundance of K and R amino acid residues in basic IDRs (Figure 2F). As expected, IDRs containing the longest K/R motif have a significantly higher average pI than IDRs with shorter K/R motifs (Figure 2H, Supplementary Figure 11C, Supplementary Table 14). Furthermore, we analyzed the hydrophobicity of these IDRs using grand average of hydropathicity (GRAVY) calculations (Figure 2I, Supplementary Figure 12, 13A, Supplementary Table 15). All datasets have hydrophilic IDRs with mean GRAVY values between −1.4 and −2.0, irrespective of the particular K/R motif, consistent with the hydrophilic nature of IDRs in general (103–105). Next, we evaluated the GRAVY distribution between the datasets and each of the three K/R motifs (Figure 2I, Supplementary Figure 13B). The RDPA proteome possesses K/R motif-containing IDRs with a significantly higher average hydrophilicity compared to other datasets, with the exception of the nucleo-specific proteome with the longest K/R motif. Furthermore, IDRs with the longest K/R motif are significantly more hydrophilic than IDRs with shorter K/R motifs, regardless of the dataset (Supplementary Figure 13C, Supplementary Table 15). We suggest that the multimodal distribution of GRAVY of the RDPA and cytosol-specific proteomes (Supplementary Figure 13B) is caused by low protein counts in these datasets.

In summary, these data show that RDPA proteins specifically contain longer IDRs with more charged amino acid residues than other datasets. This is consistent with the previous observation that PIP2 nuclear effectors associate with charged inositol headgroups, presumably via their hydrophilic protein regions (79).

### The analysis of posttranslational modifications in RDPA proteome

Charge is a crucial parameter of the components of biomolecular condensates. The formation of a condensate represents a metastable state when the charge is in a desirable balance. PTMs, such as phosphorylation, often induce the collapse of this balance, ultimately leading to the dissolution of a condensate (36,41,106–108). Therefore, we searched for sites of common PTMs, namely acetylation, methylation, phosphorylation, SUMOylation and ubiquitination, within IDRs containing K/R motifs using the PhosphoSitePlus database (82). Indeed, phosphorylation was the most abundant PTM across all datasets and the three K/R motifs analyzed, with the highest incidence in the RDPA proteome (Table 2, Supplementary Table 16). Phosphorylation is a very common PTM of IDRs, usually associated with a negative effect on the propensity for phase separation (108). It has been shown that differential phosphorylation of the intrinsically disordered C-terminal domain of Pol2 alters its condensation capacity and integration into transcription initiation and splicing condensates (41,109).

**Table 2.**
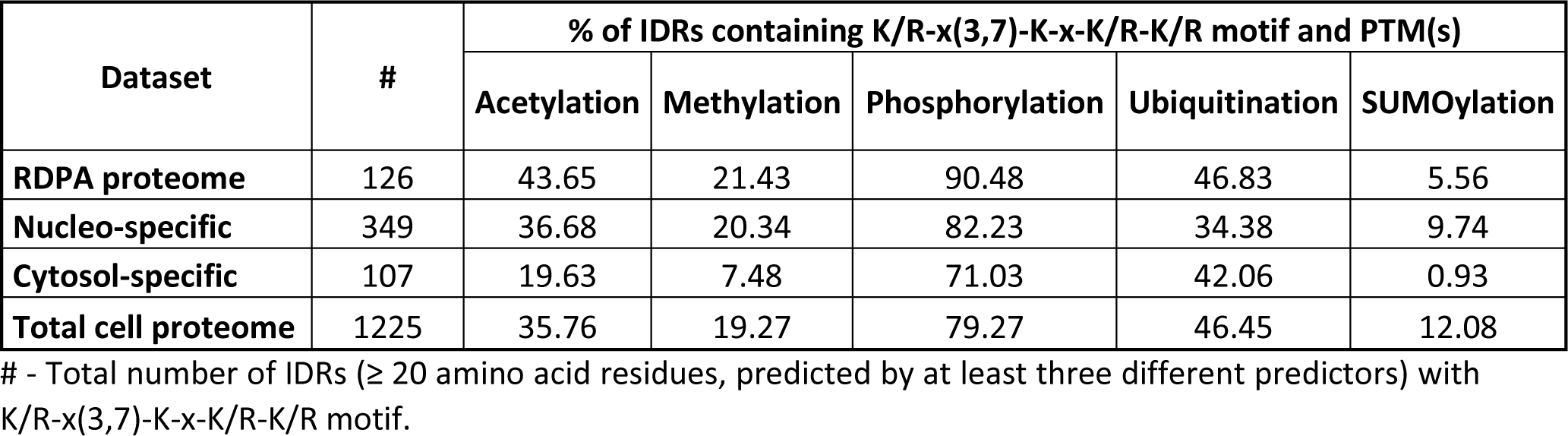
Frequency of posttranslational modification sites within IDRs containing K/R motif.

Furthermore, RDPA proteome IDRs containing the K/R-x(3,7)-K-x-K/R-K/R motif were enriched for all screened PTM sites except SUMOylation (Table 2). On the contrary, the cytosol-specific proteins generally showed a low rate of acetylation, methylation and SUMOylation PTMs, which is in agreement with the literature (110–112). Acetylation and methylation are two very common modifications of histone proteins that determine chromatin accessibility and thus the rate of transcription (113). The subcompartmentalization of differentially active chromatin is driven by the phase separation, suggesting that these PTMs are indeed critical features that define the dynamic nuclear architecture and influence gene expression (114,115). The above data show that PIP2-associated IDRs are sites of intense PTM regulation, suggesting their importance in regulating processes that depend on protein condensation capacity.

### RDPA proteins participate in the regulation of gene expression

GO analysis provides valuable insights into the function of proteins identified by shotgun MS-based approaches. Therefore, the biological processes in which RDPA proteins are involved were analyzed using Metascape (67). The results showed that RDPA proteins are mainly involved in different stages of gene expression, including chromatin accessibility, RNA transcription, RNA processing and RNA transport (Figure 3A, Supplementary Tables 17, 18). These processes, such as Pol2 transcription, RNA processing or RNA export, depend on the formation of distinct membraneless nuclear compartments in a process regulated by RNA (25,36). This observation is consistent with our bioinformatic data (Figure 2) as well as with previously published data on PIP2 effectors (2,10,79,116).

**Figure 3:**
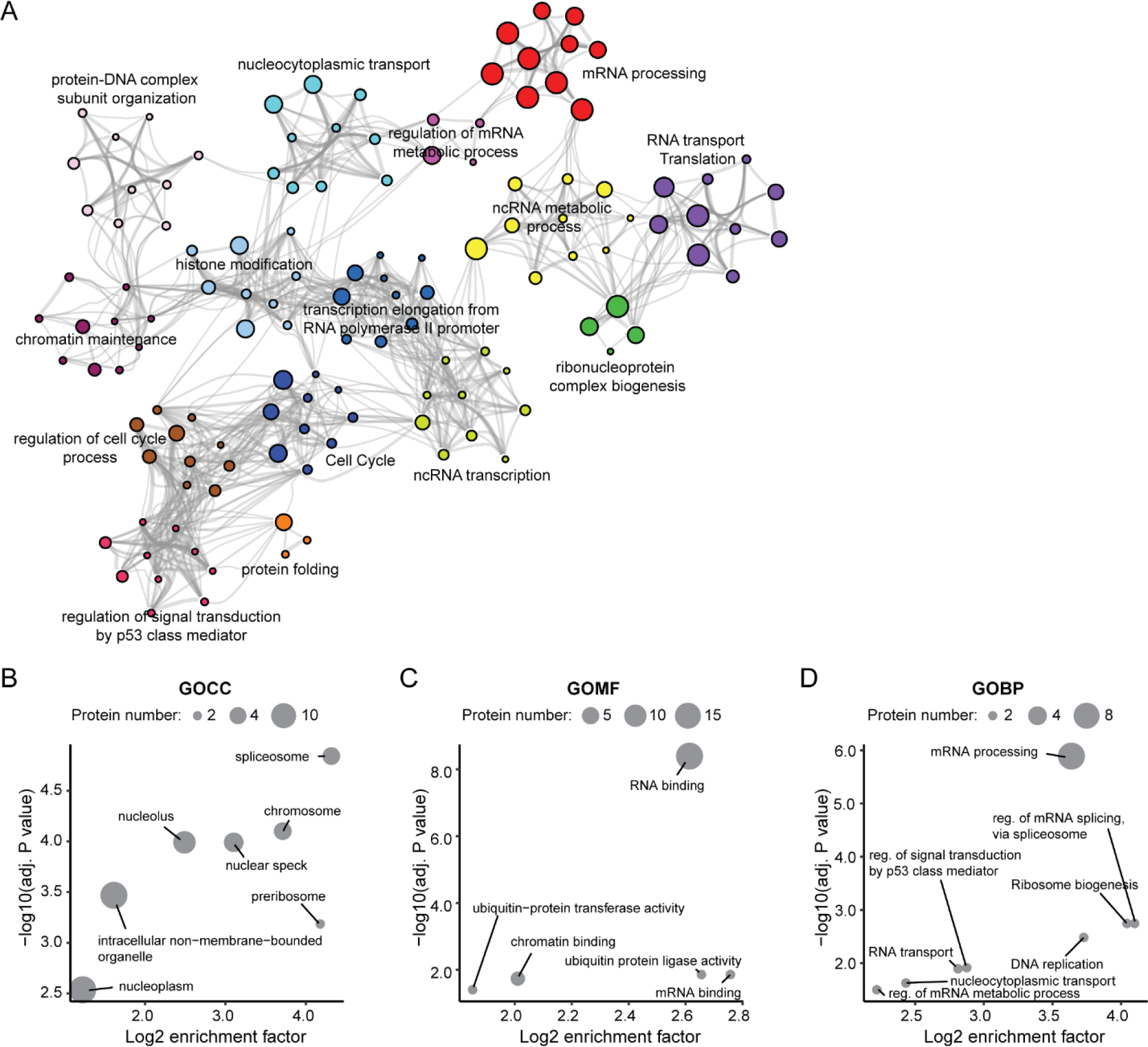
Functional analysis of the RDPA proteome. **A)** Hierarchical clustering of functional terms overrepresented in the RDPA proteome revealed enrichment of proteins regulating gene expression. Each node represents a term; the size and color represent the number of input genes and cluster identity, respectively. Terms with a similarity score > 0.3 are linked by an edge. Representative terms are selected for each cluster and shown as label. The analysis was performed in Metascape. **B-D)** Gene ontology (GO) analysis of human proteins containing K/R-x(3,7)-K-x-K/R-K/R motif in IDRs using SLiMSearch tool based on **(B)** cellular compartment (GOCC), **(C)** molecular function (GOMF), and **(D)** biological process (GOBP). The y-axis shows the –log10 adjusted p-value (Fisher’s exact test) of proteins from a GO category, the x-axis shows the log2 enrichment factor. The size of the bubble corresponds to the number of proteins.

To verify the GO results, we took a reverse approach and screened the human proteome for the presence of PIP2-binding K/R motifs within the IDRs. We used a short and linear motif discovery tool – SliMSearch (81) - with a stringent disorder score cut-off to have a high probability of the motifs being located within the IDRs. The SLiMSearch results showed that the K/R-x(3,7)-K-x-K/R-K/R motif was present in 29 proteins, the K/R-x(3,7)-K-x-K/R motif was present in 61 proteins and the K/R-x(3,7)-K-x-K motif was present in 43 proteins. We further investigated whether the K/R motifs are preferentially localized in nuclear IDR-containing proteins, as suggested by the data shown in Figure 2C. Indeed, GO localization analysis showed that all three K/R motifs are enriched in proteins associated with nuclear components such as nuclear speckles and nucleoli (Figure 3B). In addition, two shorter motifs were enriched for nuclear euchromatin and nucleosome (Supplementary Figure 14A, 15A, Supplementary Table 19, 22, 25). Thus, these data are consistent with our previous observations and suggest that K/R motifs within the IDRs have specific roles in nuclear processes.

We further focused on elucidating the molecular functions of these proteins (Figure 3C, Supplementary Figure 14B, 15B, Supplementary Table 20, 23, 26). Our analysis confirmed that proteins with at least one of the three K/R motifs in the IDRs have RNA binding capacity. These data are consistent with our bioinformatic analysis, which found that RNA binding function is enriched in the RDPA proteome (Figure 2A). The RNA binding ability of nuclear proteins is an important feature for the formation of biomolecular condensates, supporting the notion that RNA is the key factor defining nuclear compartmentalization (86). Next, we analyzed the biological processes of proteins with K/R motifs containing IDRs. Processes of RNA splicing, RNA transport, ribosome biogenesis, nucleocytoplasmic transport and regulation of signal transduction by p53 class mediator were enriched between all motifs (Figure 3D, Supplementary Table 21, 24, 27). In contrast, histone modifications, nucleosome positioning, transcription elongation from the RNA polymerase II promoter and ncRNA metabolism were enriched specifically for proteins with shorter K/R motifs in IDRs (Supplementary Figure 14C, 15C, Supplementary Table 24, 27).

Taken together, these results are consistent with the results of the RDPA proteome bioinformatics and GO analyses (Figure 2A, C and Figure 3A). Interestingly, SLiMSearch data suggest that proteins with K/R-x(3,7)-K-x-K/R-K/R motif in IDRs may have a different nuclear distribution and be involved in different processes than proteins with IDRs with shorter K/R motifs, indicating a degree of specificity for a particular site of action. We hypothesize that different K/R motifs may localize to different PIP2 nuclear sub-populations (nuclear speckles, nucleoli and NLIs) and thus PIP2 acts as a molecular wedge via RNA association to attract and retain different sets of RDPA proteins. The PIP2-dependent landscape of subnuclear localization has recently been suggested as an important determinant of nuclear architecture (92). A similar dependence on K/R protein levels has been identified as an important determinant of protein phase separation and localization (29,117). Furthermore, a recently published study shows that mutations in IDR-containing proteins are often associated with the formation of K/R frame shifts that alter their phase separation capacities, leading to progression of cancer predispositions (30). Thus, we propose that PIP2 represents a novel player that affects nuclear compartmentalization through phase separation of interacting proteins.

### RNA regulates the association of the RDPA protein BRD4 with PIP2

Our bioinformatic analyses revealed that RDPA proteins contain charged K/R motifs within their IDRs, which are thought to be responsible for PIP2 recognition (2,79). The RDPA protein BRD4 is an IDR-containing transcriptional regulator with the ability to form phase-separated condensates *in vitro* and *in vivo* (118). To characterize the interactions between the selected RDPA protein BRD4 and PIP2, we performed pull-down experiments using PIP2-conjugated beads to mimic naturally occurring PIP2 structures under different conditions. Our results showed that protein BRD4 associates with PIP2 structures in a RNA-dependent manner, as the addition of exogenous RNA positively affected its PIP2 binding capacity (Figure 4A). Furthermore, the interactions between PIP2 and BRD4 protein is of electrostatic nature, as increased NaCl concentration (300 mM) significantly decreased PIP2 binding (Figure 4A). Biomolecular condensates are sensitive to increasing salt concentration because salt ions reduce weak electrostatic interactions necessary for cohesive forces within condensates (101,119). NH_4_OAc treatment prevents the formation of structured RNA folds by denaturing RNA molecules, while leaving protein structures intact (36). This treatment revealed that RNA-RNA interactions and the formation of higher order folds (i.e., dsRNA) are important prerequisites for PIP2-BRD4 protein interactions (Figure 4A). The 1,6-hexanediol is often used to dissolve protein condensates formed by phase separation *in vitro* and *in vivo* (120), although the reliability of its use *in vivo* is still the subject of intense debate. Recent studies have shown that 1,6-hexanediol may affect the chromatin state and enzymatic functions of some proteins (121,122), making 1,6-hexanediol the drug of choice for *in vitro* assays only. The 1,6-hexanediol in the pull-down reaction reduced PIP2 association with the BRD4 protein (Figure 4A). The addition of the crowding agent dextran partly restored the association of PIP2 with the BRD4 protein under conditions where no RNA was added (Figure 4A). Similar results were obtained with PIP2-containing nuclear material pulled down through the GST-PLCδ1 PH domain, suggesting the relevance of these observations to naturally occurring PIP2-BRD4 complexes (Figure 4B).

**Figure 4:**
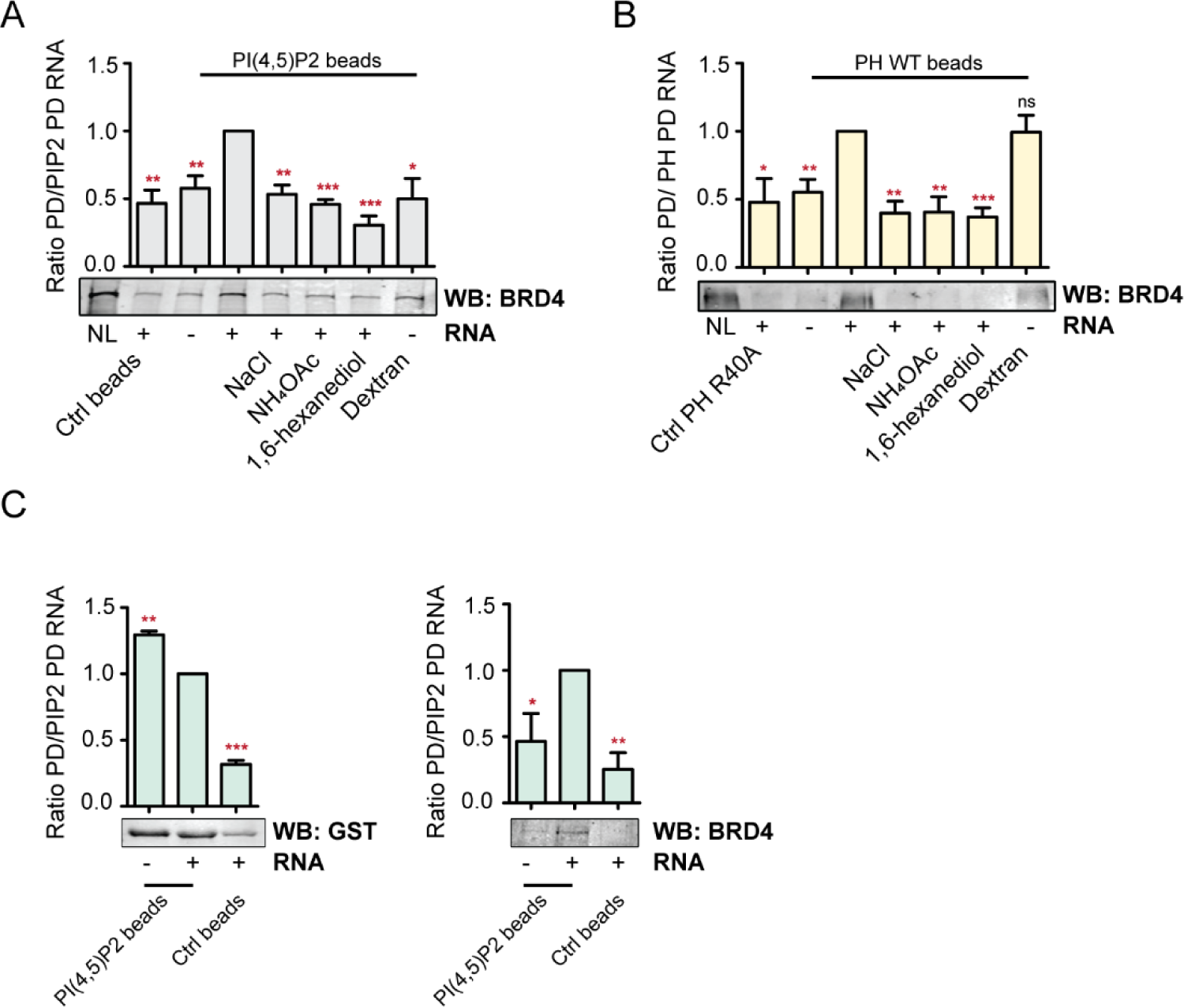
PIP2-conjugated agarose beads pull-down assays from nuclear lysates with added nuclear RNA extract upon different conditions. The PIP2 and empty control beads were incubated for 1 h at 4 °C in nuclear lysates, washed, and subjected to WB detection of BDR4 protein **(A)**. WB signals at each pull-down condition in every repetition were normalized to the highest signal. Statistical analysis was performed using Student’s t-test (n = 4). **(B)** GST-PLCδ1 PH pull-down assay testing the significance of PIP2 pull-down results for naturally occurring PIP2-BRD4 structures. WB signals at each pull-down condition in every repetition were normalized to the highest signal (GST-PLCδ1 PH WT domain + RNA). Statistical analysis was performed using Student’s t-test (n = 6). **(C)** PIP2-conjugated beads pull-down assay with spike-in of recombinant GST-PLCδ1 PH domain testing the specificity of the effect of RNA on PIP2 binding of BRD4 protein. WB signals at each pull-down condition in every repetition were normalized to the highest signal of PIP2-beads upon RNA extract addition. Statistical analysis was performed using Student’s t-test (n = 4). NL – nuclear lysate, * P < 0.05, ** P < 0.001, *** P < 0.0005

To further test the specific effect of RNA on BRD4 protein, we set up an experiment where we added the same amount of purified recombinant GST-tagged GST-PLCδ1 PH domain to the nuclear lysate as an internal control. We compared the effect of RNA extract addition on PIP2 association with BRD4 protein and the control PH domain (Figure 4C). The addition of RNA extract increased the PIP2 association of BRD4 protein, whereas the association of the GST-PLCδ1 PH domain with PIP2 was decreased under these conditions. These results suggest that the same amount of RNA that stimulates PIP2 association with K/R IDR-containing BRD4 proteins negatively affects PIP2 recognition by the canonical PIPs-binding PH domain. Thus, RNA appears to act as a local scaffold rather than a non-specific crowding agent. Based on these data, we hypothesize that the structured RNA attracts the specific RNA-binding proteins and thus locally increases their concentration, eventually leading to the formation of a condensate in the vicinity of a PIP2-containing surface.

### BRD4 protein foci localize non-randomly in the vicinity to PIP2-containing nuclear structures

Nuclear PIP2 localizes to transcriptionally relevant compartments - nuclear speckles, NLIs and nucleoli (7). Therefore, we investigated whether the transcriptional regulator BRD4 protein localizes to nuclear PIP2 structures. To visualize the detailed localization of BRD4, we used super-resolution microscopy, which has recently been successfully applied to assess co-patterning of nuclear proteins with PIP2 (92). Our data showed that BRD4 formed foci in a dispersed pattern inside the nucleus (Figure 5A). We analyzed the colocalization of the signal of BRD4 protein with the PIP2 signal and compared the real data with the data randomized by rotating one channel with respect to the second channel (57). The following coefficients were used to test signal colocalization and spatial codistribution: Pearson’s coefficient, which measures the degree of correlative variation between the two channels. Manders’ M1 and M2 coefficients, which measure the proportion of the intensity in each channel that coincides with an intensity in the other channel. The M1 and M2 coefficients are less dependent on the actual intensity ratios between the channels. Spearman’s coefficient, which detects the mutual dependencies of two channel signals and thus measures the statistical association between these two channel signals. Statistical analysis of the data revealed that BRD4 protein colocalized to a very limited extent, but rather tended to localize in the vicinity of PIP2 (Figure 5B).

**Figure 5:**
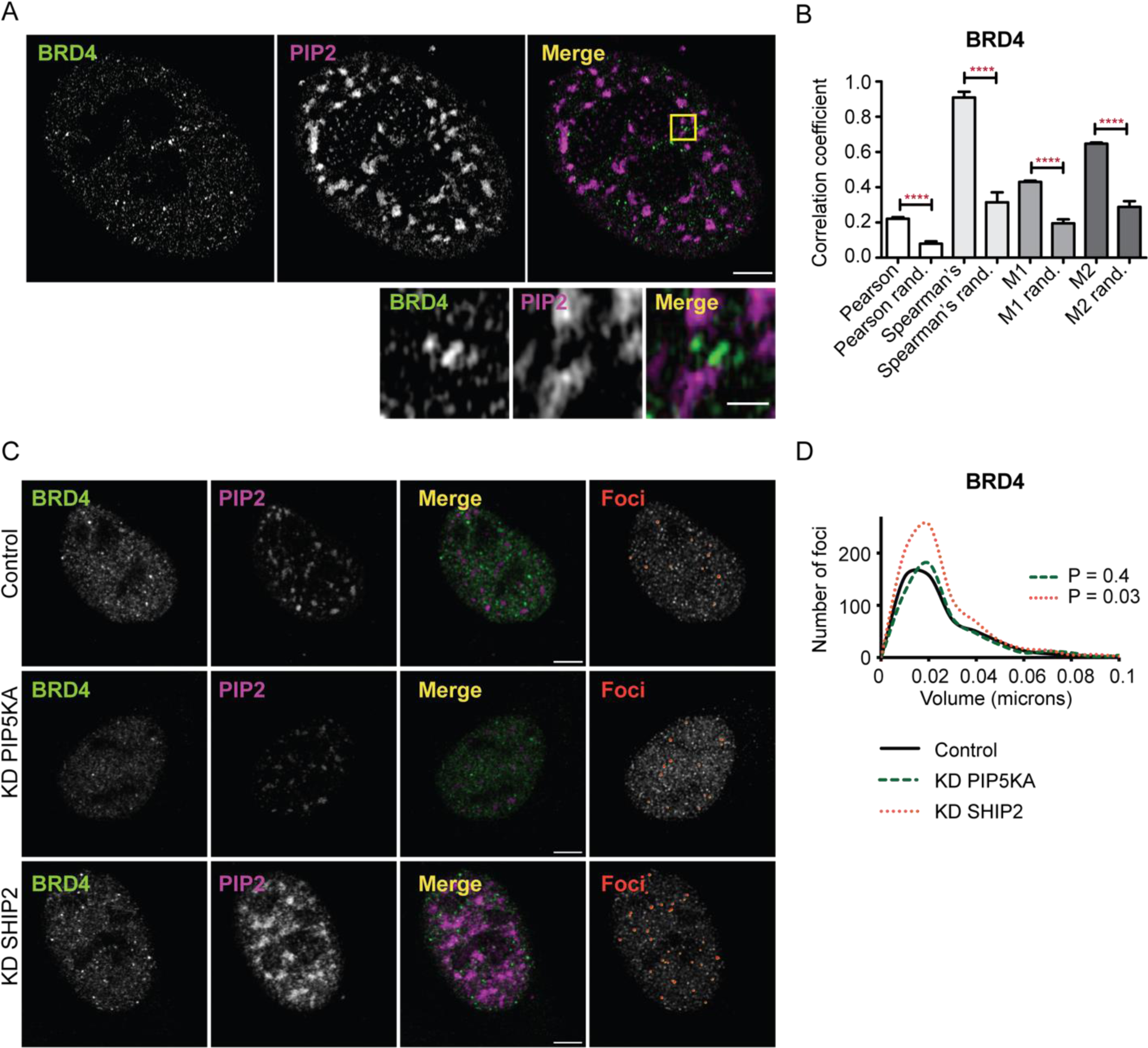
Colocalization of BRD4 with PIP2 and changes in BRD4 nuclear localization induced by the manipulation of PIP2 levels visualized by super-resolution and confocal microscopy. **(A)** Representative images of immunofluorescence staining for BRD4 and PIP2 show the localization of this protein in the vicinity of PIP2 in the nuclei of U2OS cells. The image acquisition was performed by structured illumination microscopy (SIM). The inset shows a detail of BRD4 and PIP2 localization at a nucleoplasmic region within the U2OS cell nucleus (yellow square). Scale bars correspond to 5 µm and 1 µm resp. (inset). **B)** Statistical analysis of colocalization parameters by Pearson’s, Spearman’s, and Manders’ coefficients M1 and M2 compared to randomized images was performed using Student’s t-tests. Error bars correspond to SEM (** P < 0.005, *** P < 0.001, **** P < 0.0001), n = 3, N = 34 cells. **(C)** U2OS cells with decreased PIP2 levels by depletion of PIP5KA and increased PIP2 levels by depletion of SHIP2, respectively. Representative figures show the localization of PIP2 and BRD4 using immunofluorescence staining. The last column shows the identified foci in 3D of representative images in red. Scale bars correspond to 5 µm. **D)** The chart visualizes the number of foci and the volume in µm^3^ of identified foci per BRD4 in control (solid line), knockdown (KD) PIP5KA (green dashed line), and KD SHIP2 (orange dotted line) U2OS cells. Frequencies distributions were analyzed using the Kolmogorov-Smirnov test (n = 3, control N = 26, KD PIP5KA N = 26, and KD SHIP2 N = 33 cells, respectively).

Hence, it seems that BRD4 protein foci do not directly colocalize with PIP2 but instead exhibit non-random co-clustering, suggesting an indirect interaction. This implies a potential association with the surface of nuclear PIP2 structures via other molecules, as hinted by the findings presented in Figure 4A, which propose structured RNA involvement.

### Higher PIP2 levels increase the number of nuclear BRD4 protein foci

Based on the data presented so far, we propose that nuclear PIP2 attracts RDPA proteins containing K/R motifs, leading to their localization in PIP2-enriched areas. Therefore, the localization patterns of BRD4 protein should vary under conditions of different nuclear PIP2 levels. To test the effect of varying PIP2 levels on the localization of BRD4 protein, we experimentally increased nuclear PIP2 levels by depleting SHIP2 phosphatase and conversely decreased PIP2 levels by depleting PIP5KA (Supplementary Figure 16A).

Image analysis and quantification of the microscopy data showed that increased PIP2 levels increased the number of BRD4 nuclear foci (Figure 5C, 5D). This effect may help to explain the importance of nuclear PIP2 in transcription and Pol2 condensation described elsewhere (7,10). The total protein level of BRD4 was slightly lowered upon SHIP2 depletion, suggesting that increase in BRD4 protein foci number is not due to increased total protein concentrations, but rather due to changes in its local level (Supplementary Figure 16B.

These data suggest that the level of nuclear PIP2 does indeed influence the formation or stability of BRD4 protein foci, presumably by altering its local concentration via recruitment. The presence of an amphiphilic molecule in the nuclear environment has been suggested to explain the formation of so-called microemulsions, which are typically organized in nuclear processes (125). Based on our data, we propose that nuclear PIP2 regulates the affinity between RNA and BRD4 protein. This interaction define the areas of foci formation, thus orchestrating nuclear subcompartmentalization and leading to efficient gene expression. Our data are consistent with the recently published work in which the authors describe, among other fundamental findings, the effect of PIPs on the quantity, size and morphology of condensates in an *in vitro* system (Dumelie et al., 2023). This model brings new perspectives to another recent observation that carcinogenic human papillomavirus (HPV) infection increases nuclear PIP2 levels in human wart samples (93). Thus, we provide mechanistic insights into the real biological implications of this phenomenon.

In conclusion, we have shown that nuclear PIP2 localization is dependent on the presence of dsRNA and identified RDPA proteins that associate with nuclear PIP2 in a dsRNA-dependent manner. To this end, we developed and optimized a MS-based experimental pipeline that can be applied to other nuclear lipid-protein interactors. Our results showed that interactions between RDPA protein BRD4 and PIP2 are mediated by electrostatic interactions, presumably via the enriched K/R motifs within the IDRs. The PIP2-binding function of such K/R motifs has been described previously (3,79). Here, we have shown that more than half of the RDPA proteins have experimentally demonstrated RNA binding capacity and are associated with the process of phase separation. We found that IDRs have a bimodal distribution of pI and we provide bioinformatic tools for their convenient analyses. We also showed that IDRs of RDPA proteins are longer and more hydrophilic. The K/R motif-containing IDRs of RDPA proteins are enriched for phosphorylation, acetylation and ubiquitination sites, providing a further level of regulation for their integration into phase-separated condensates. GO analysis revealed that RDPA proteins are involved in RNA transcription, RNA processing and transport, translation, cell cycle regulation, histone modification and chromatin maintenance. The involvement of proteins containing IDRs with K/R motifs in similar processes was confirmed by our SLIMsearch analysis. We showed the localization of RDPA protein BRD4 with respect to nuclear PIP2 structures. Finally, we have shown that increased levels of nuclear PIP2 leads to increased number of BRD4 foci.

## Acknowledgments

We acknowledge Mgr. Karel Harant and Mgr. Pavel Talacko from Laboratory of Mass Spectrometry, Biocev, Charles University, Faculty of Science for performing LC-MS/MS measurements. We are grateful to Dr. Ivan Novotný from light microscopy facility at the Institute of Molecular Genetics for his technical support in super resolution microscopy, Dr. Michaela Blažíková and Mgr. Jan Valečka for their help with microscopy data analysis. Pavel Kříž and Iva Jelínková for technical support and Dr. Helena Kupcová Skalníková for proofreading of the manuscript.

## Funding

This work was supported by the Grant Agency of the Czech Republic (Grant nos. 19-05608S and 18-19714S); by the project National Institute for Cancer Research (Programme EXCELES, ID Project No. LX22NPO5102) by the European Union - Next Generation EU; by the Czech Academy of Sciences (Grant no. JSPS-20-06); by the Institutional Research Concept of the Institute of Molecular Genetics (Grant no. RVO: 68378050) and of the Institute of Animal Physiology and Genetics (Grant no. RVO: 67985904); by the MEYS CR (COST Inter-excellence internship LTC19048 and LTC20024) and the project: „BIOCEV – Biotechnology and Biomedicine Centre of the Academy of Sciences and Charles University” (CZ.1.05/1.1.00/02.0109), from the European Regional Development Fund. This publication is based upon work from COST Action 19105-Pan-European Network in Lipidomics and EpiLipidomics (EpiLipidNET) supported by COST (European Cooperation in Science and Technology). The Microscopy Centre was supported by the MEYS CR (LM2018129 and LM2023050 Czech-BioImaging) and by the European Regional Development Fund-Projects (no. CZ.02.1.01/0.0/0.0/16_013/0001775 and CZ.02.1.01/0.0/0.0/18_046/0016045).

## Conflict of interest

The authors declare no conflict of interest.

## Notes

### Competing Interest Statement

The authors have declared no competing interest.

