## supplementary figures for "The RNA-dependent interactions of phosphatidylinositol 4,5-bisphosphate with intrinsically disordered proteins contribute to nuclear compartmentalization"

**Supplementary Figure 1**

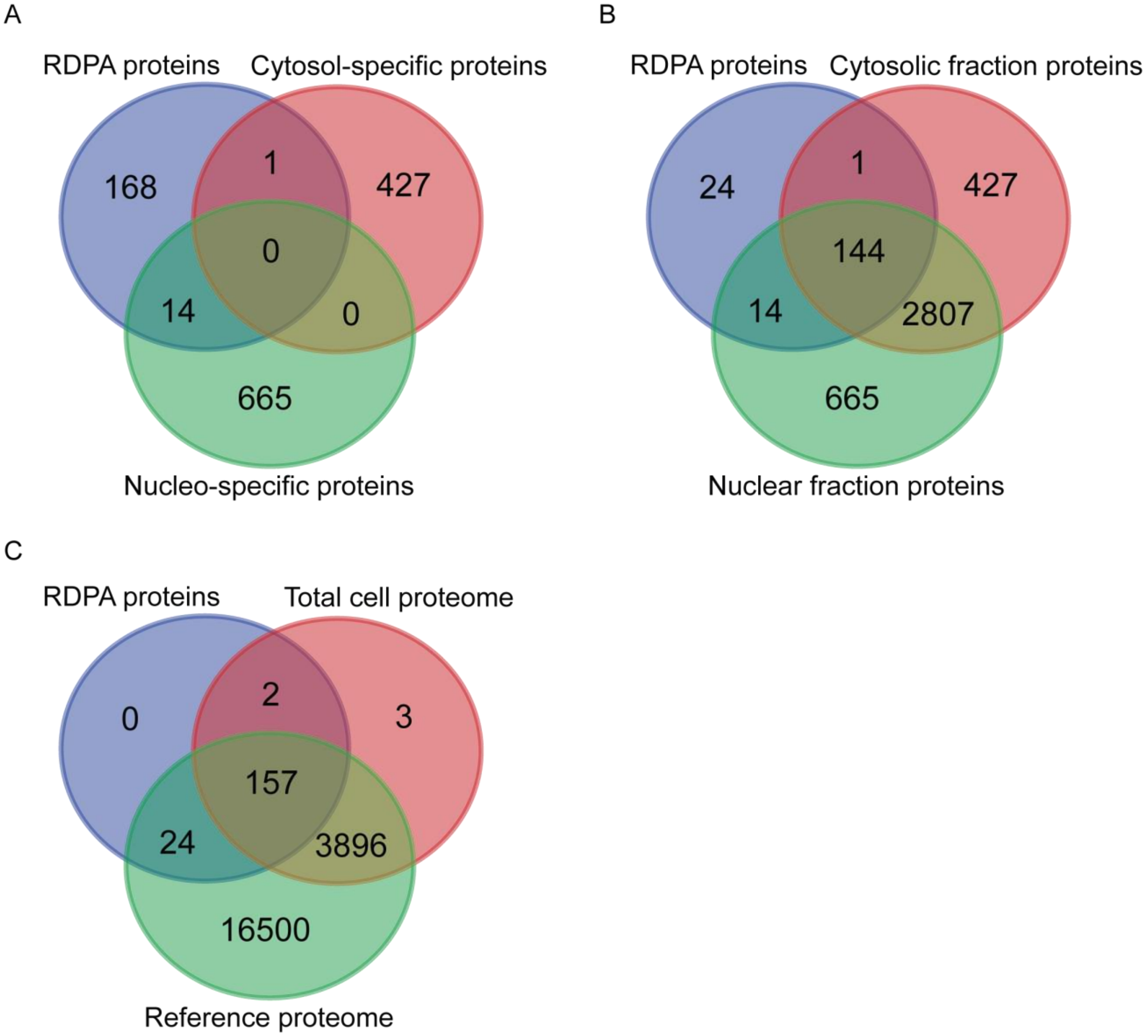


**Supplementary Figure 1: Overview and overlaps between the datasets as identified by mass spectrometry analyses (A-C).** Numbers represent proteins from Majority protein IDs mapped to UniProt (release 2022_01). Datasets Nucleo-specific proteins **(A)**, Nuclear fraction proteins **(B)** and Total cell proteome **(C)** were supplemented by missing RDPA proteins (for more information see Supplementary Table 3 and 4). The modified datasets were then used for the bioinformatic analyses presented in this study (related to Figure 2).

**Supplementary Figure 2**

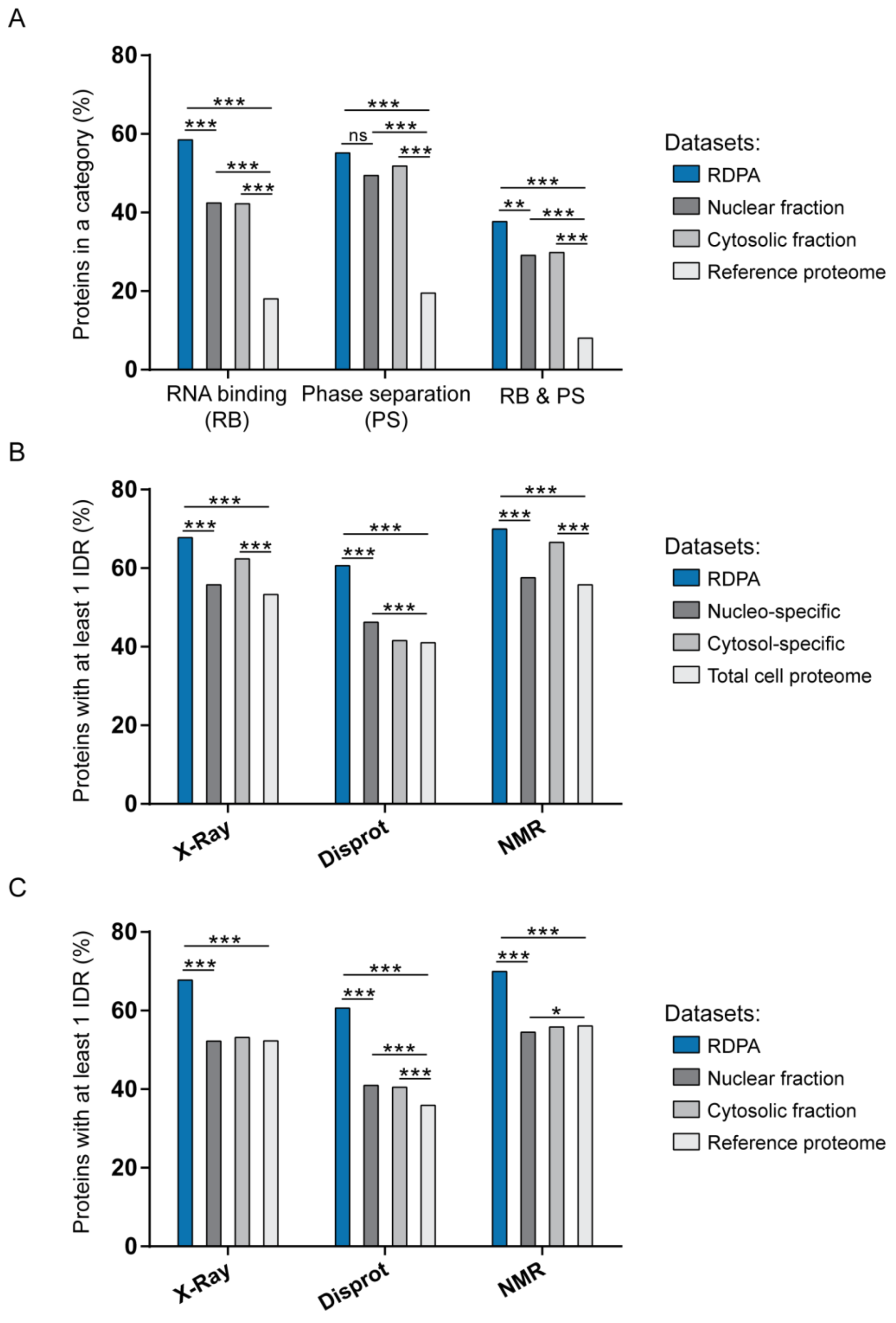


**Supplementary Figure 2: Additional bioinformatic analyses of RDPA proteome features (related to Figure 2A and B)**. **A)** RDPA proteome is significantly enriched for RNA-binding, phase separation capacity, and combination of both properties. **B-C)** RDPA proteome is enriched for IDRs longer than 30 amino acid residues predicted by ESpritz

X-Ray (X-Ray), ESpritz Disprot (Disprot), and ESpritz NMR (NMR). Statistical analysis was performed using a hypergeometric test (ns not significant, * $P < 0.05$, ** $P < 0.01$, and *** $P < 0.001$).

**Supplementary Figure 3**

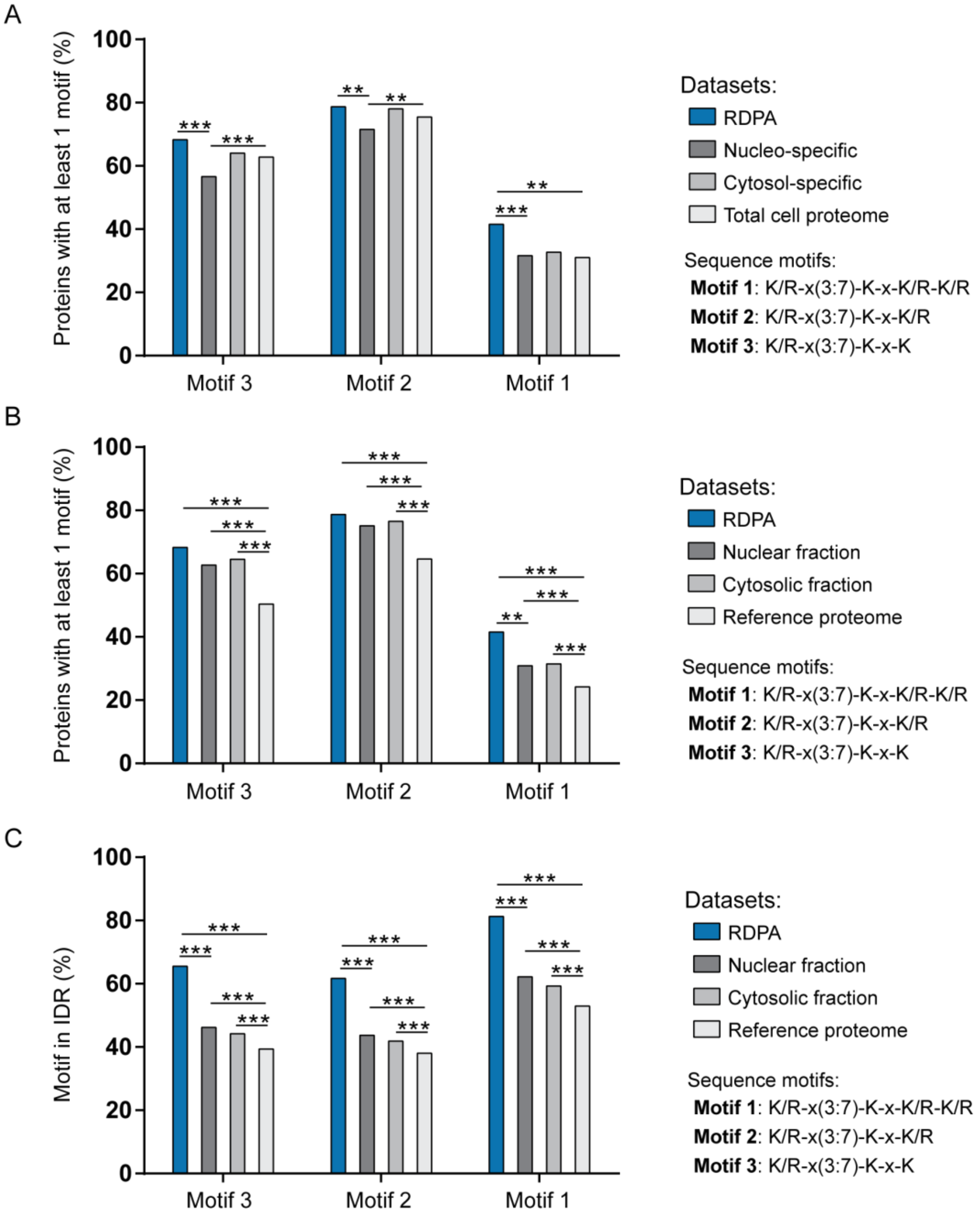


**Supplementary Figure 3: Additional bioinformatic analyses of RDPA proteome features (related to Figure 2C)**. **A-B)** Enrichment of K/R motifs in the RDPA proteome. These motifs were abundantly present in the RDPA proteome, but only the K/R-x(3,7)-K-x-K/R-K/R motif (the longest one) was significantly enriched, compared to all other datasets. **C)** Percentage of PIP2-binding K/R motif sites localized in IDRs (from all K/R motif sites in the dataset) is elevated in RDPA proteome (only IDRs predicted by at least three different predictors with minimal length 20 amino acid residues were considered). Statistical analysis was performed using a hypergeometric test (* $P < 0.05$, ** $P < 0.01$, and *** $P < 0.001$).

**Supplementary Figure 4**

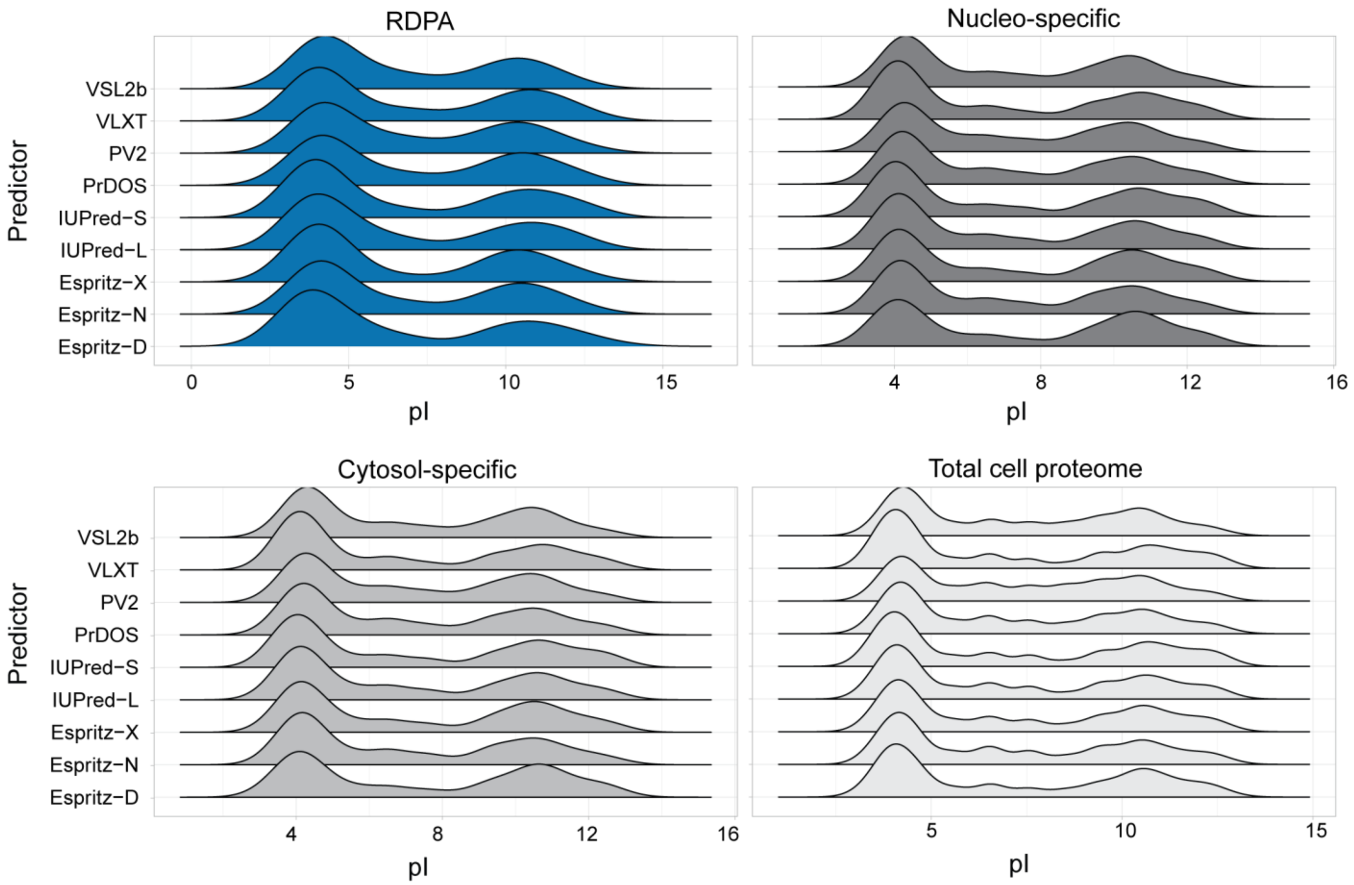


**Supplementary Figure 4: Additional bioinformatic analysis of RDPA proteome features (relevant to Figure 2D).** Bimodal pI distribution of IDRs predicted by nine different IDR predictors (Database of Disordered Protein Predictions - https://d2p2.pro/). Only IDRs with minimal length of 20 amino acid residues were considered.

**Supplementary Figure 5**

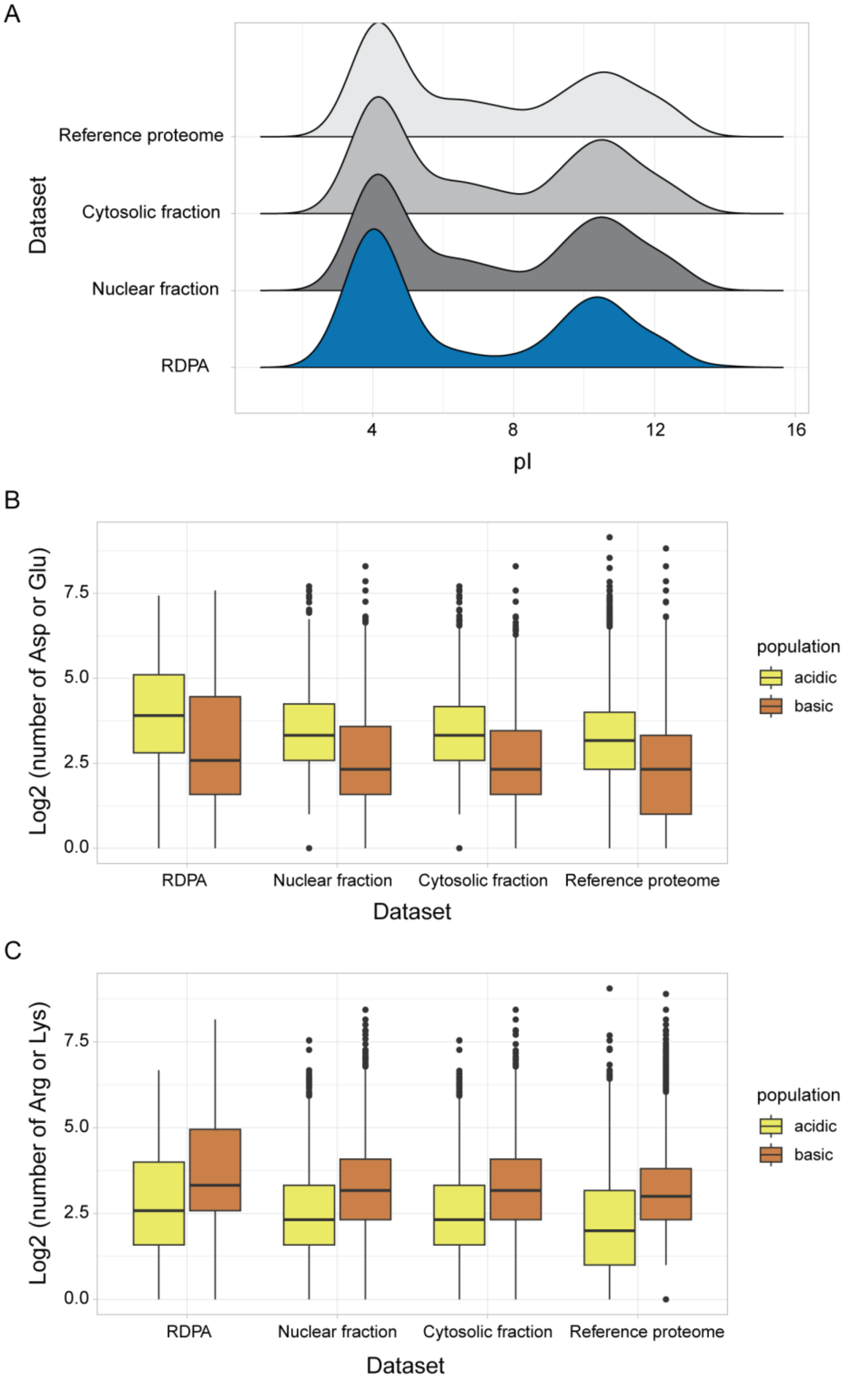


**Supplementary Figure 5: Additional bioinformatic analyses of RDPA proteome features (relevant to Figure 2D-F). A)** pI values of IDRs predicted by ESpritz X-Ray in the analyzed protein datasets show a bimodal distribution. **B)** The IDRs in the "acidic" population (pI < 7) are enriched with D/E amino acid residues. **C)** The IDRs in the "basic" population (pI > 7) are K/R-rich**.**

**Supplementary Figure 6**

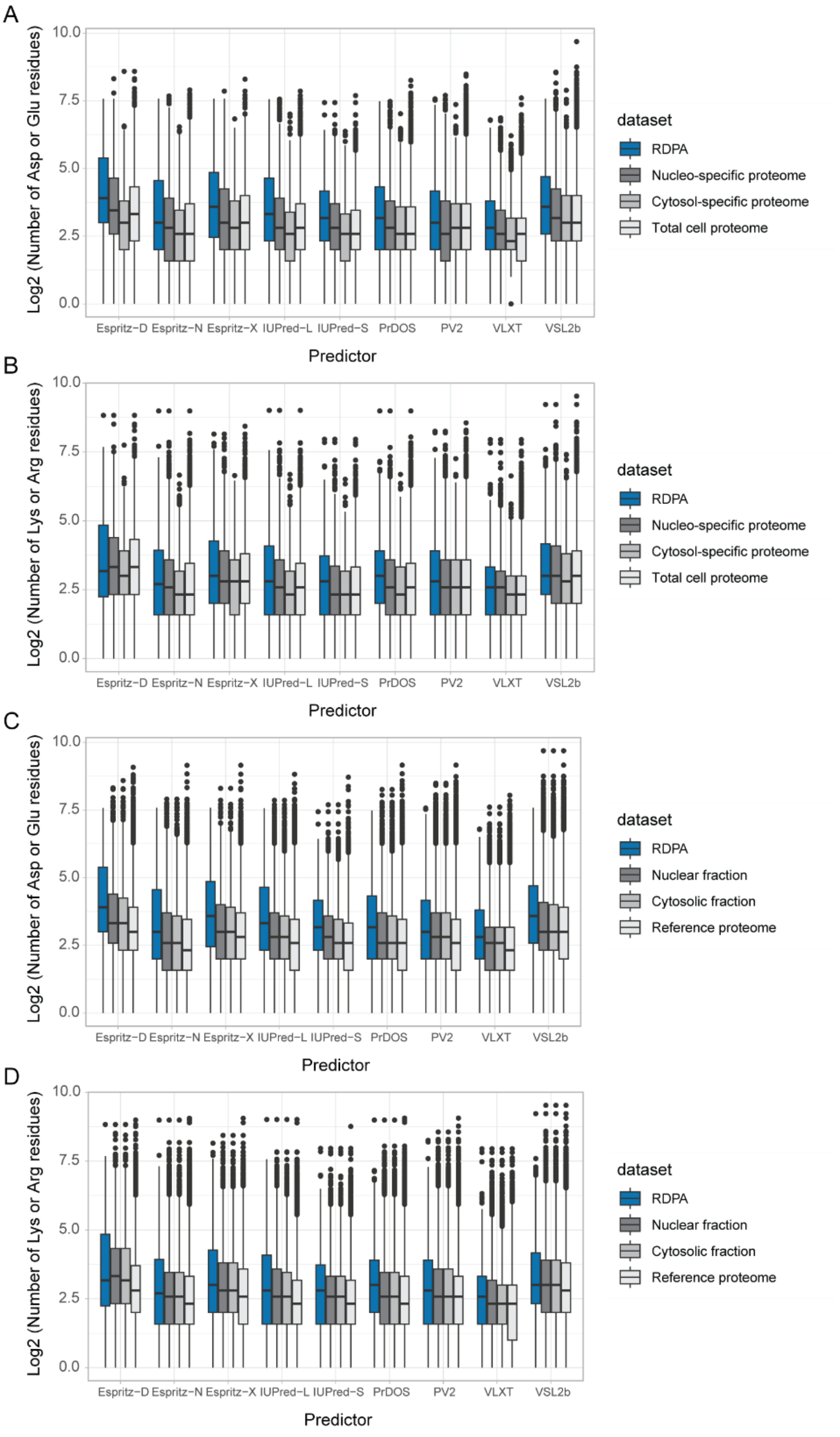


**Supplementary Figure 6: Additional bioinformatic analyses of RDPA proteome features (relevant to Figure 2E-F). A-B)** Distribution of the numbers of acidic **(A)** and basic **(B)** residues in IDRs predicted by nine different IDR predictors (Database of Disordered Protein Predictions; only IDRs with minimal length of 20 amino acid residues were considered) in the “main” datasets. **C-D)** Distribution of the numbers of acidic **(C)** and basic **(D)** residues in IDRs predicted by nine different IDR predictors (Database of Disordered Protein Predictions; only IDRs with minimal length of 20 amino acid residues were considered) in the “additional” datasets.

**Supplementary Figure 7**

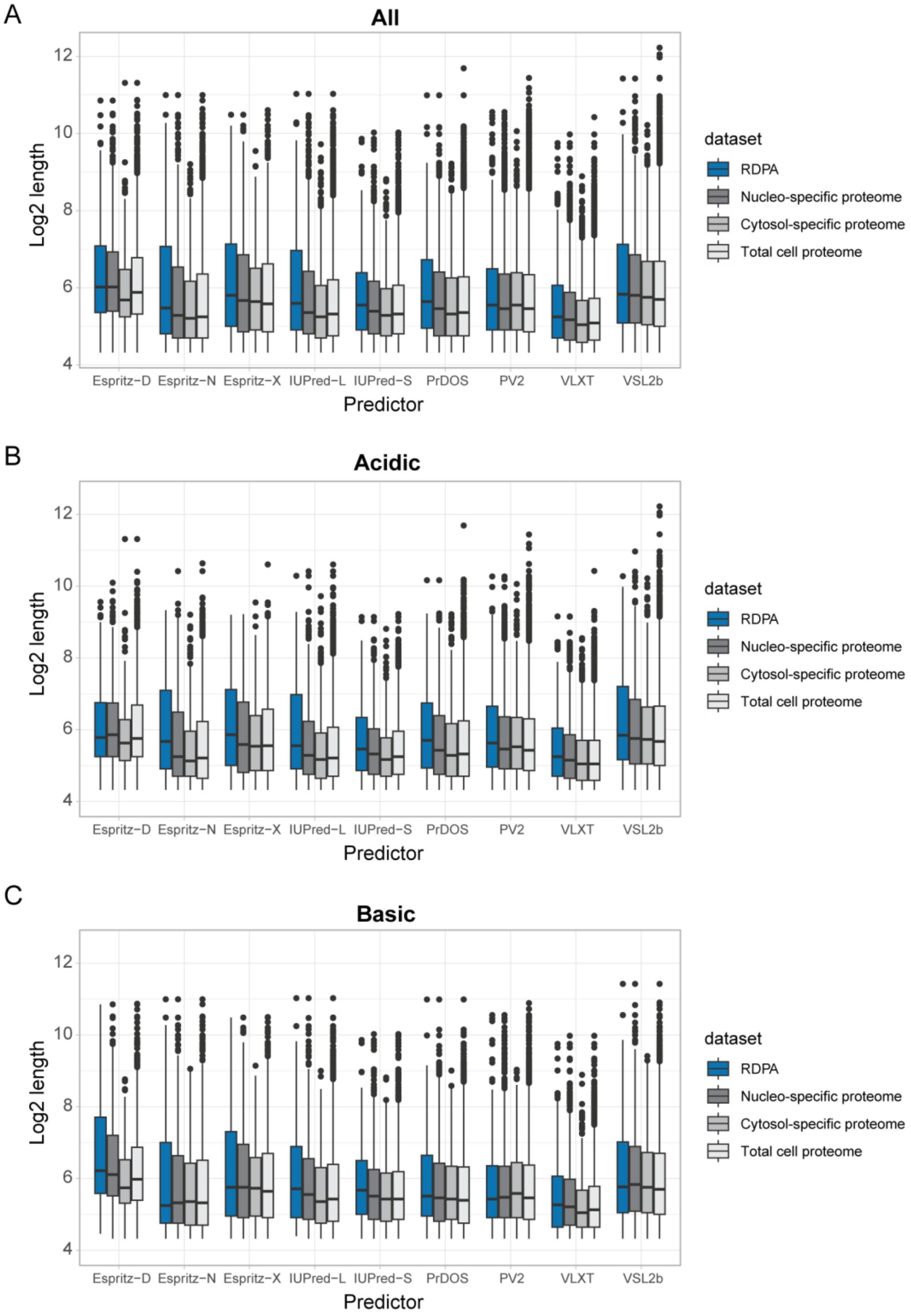


**Supplementary Figure 7: Additional bioinformatic analysis of RDPA proteome features (relevant to Figure 2D, G). A-C)** Distribution of the log2 transformed length of all IDRs **(A)** or IDRs that were acidic (pI < 7) **(B)** or basic (pI > 7) **(C)** and predicted by nine different IDR predictors (Database of Disordered Protein Predictions; only IDRs with minimal length of 20 amino acid residues were considered) in the “main” datasets.

**Supplementary Figure 8**

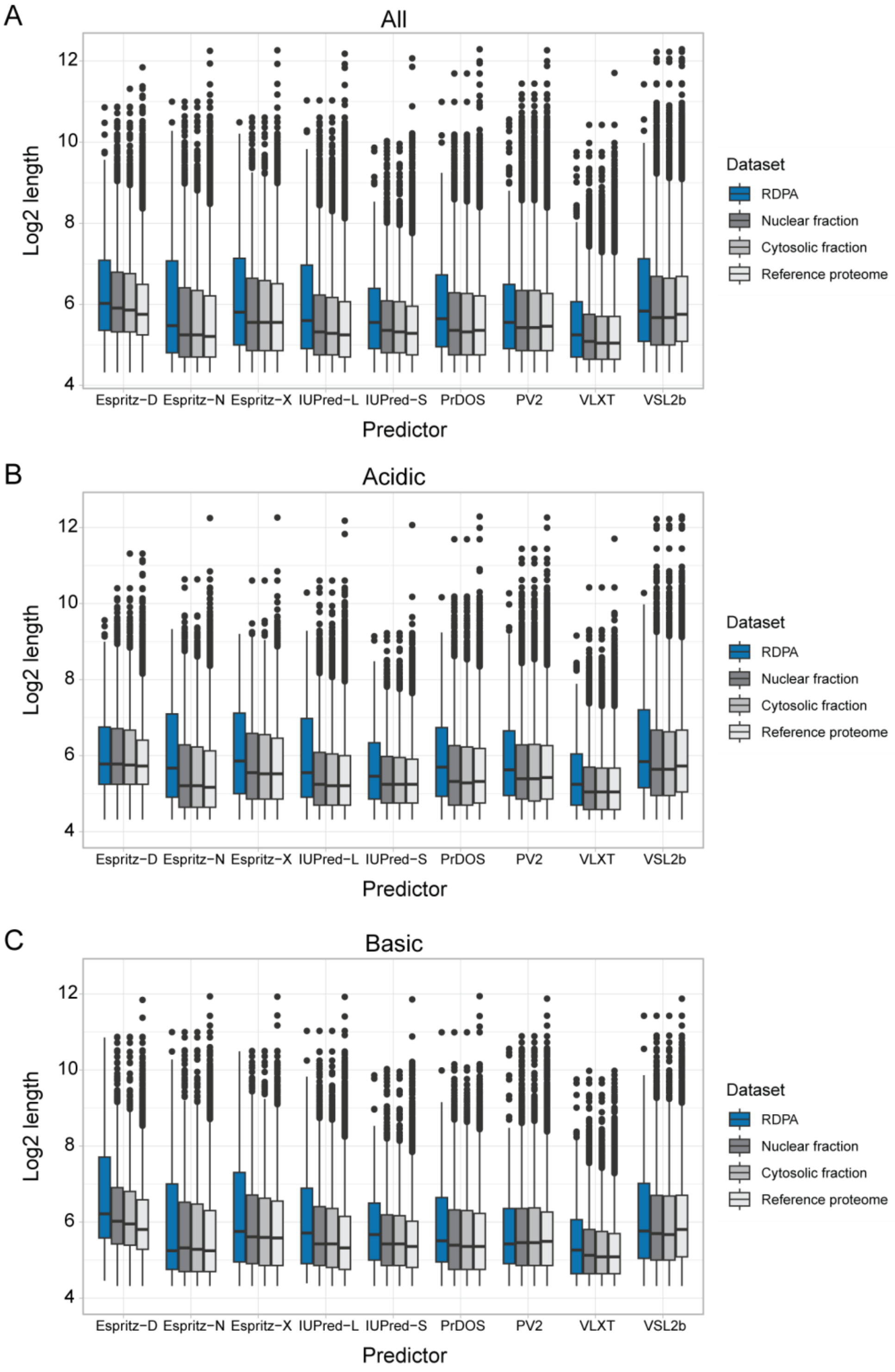


**Supplementary Figure 8: Additional bioinformatic analysis of RDPA proteome features (relevant to Figure 2D, G). A-C)** Distribution of the log2 transformed length of all IDRs **(A)** or IDRs that were acidic (pI < 7) **(A)** or basic (pI > 7) **(C)** and predicted by nine different IDR predictors (Database of Disordered Protein Predictions; only IDRs with minimal length of 20 amino acid residues were considered) in the “additional” datasets.

**Supplementary Figure 9**

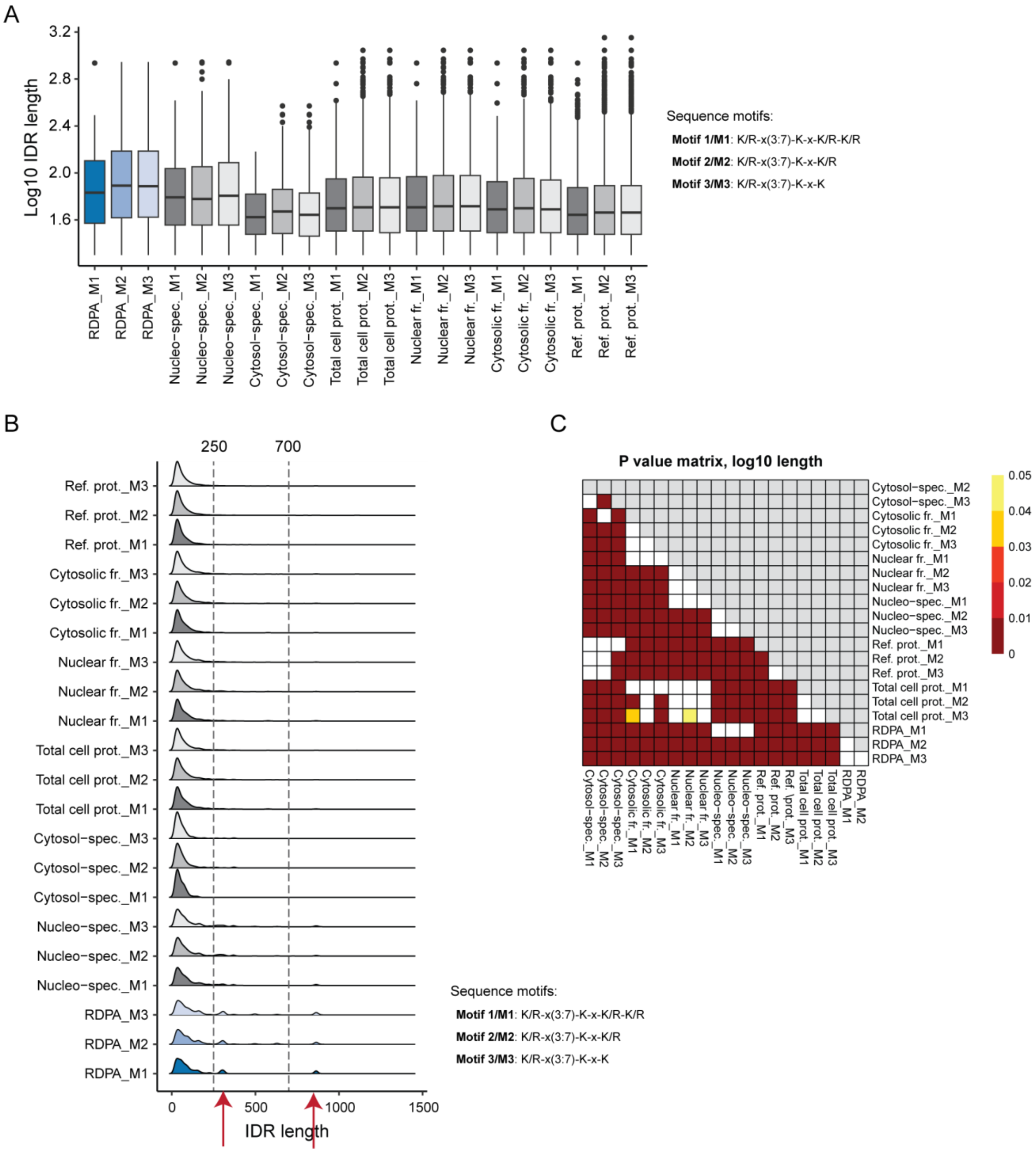


**Supplementary Figure 9: Additional bioinformatic analysis of RDPA proteome features (relevant to Figure 2G). A-C)** RDPA proteins IDRs containing the three K/R motifs tend to be significantly longer compared to the other six datasets. **A)** Boxplots show the distributions of log10 transformed length. **B)** Density plots of the IDR lengths highlight the presence of two populations of longer IDRs in the RDPA proteome and Nucleo-specific proteins (red arrows). **C)** The P values of all pairwise comparisons between the datasets and motifs were estimated by a pairwise

Wilcox test. Benjamini-Hochberg correction was applied to correct for multiple hypothesis testing. Ref. – reference, prot. – proteome, fr. – fraction, spec. – specific.

**Supplementary Figure 10**

A

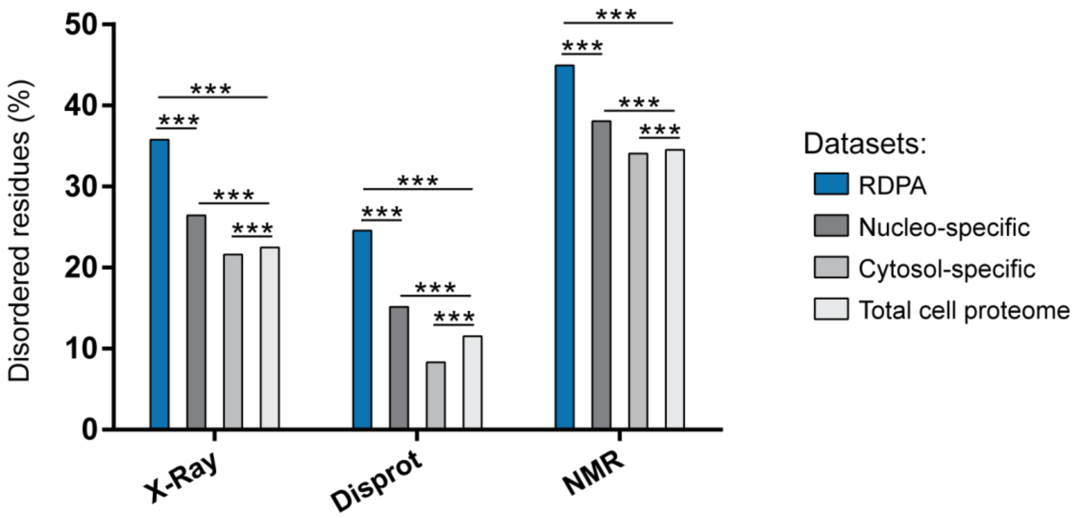


B

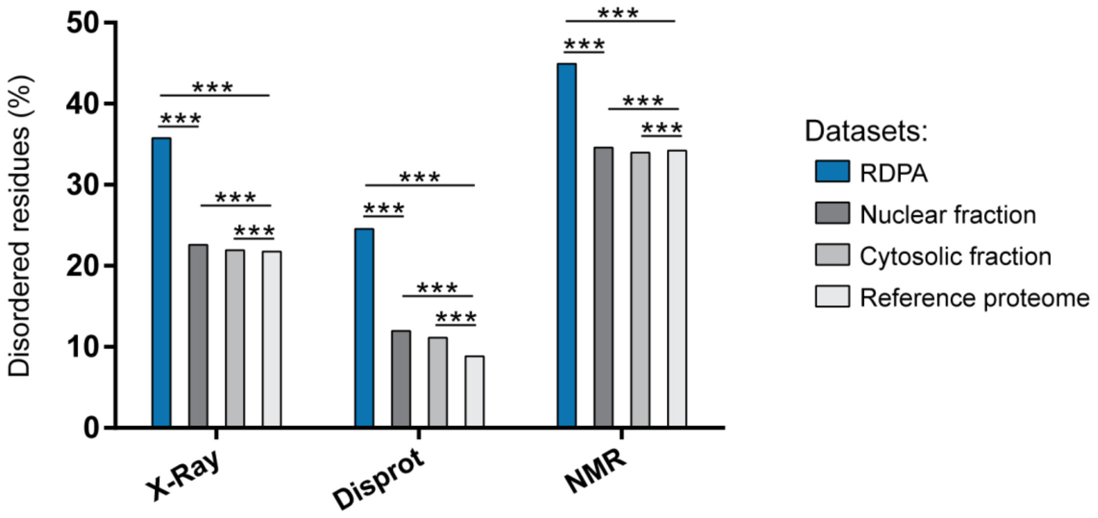


**Supplementary Figure 10: Additional bioinformatic analysis of RDPA proteome features (relevant to Figure 2B, G).** The percentage of disordered amino acid residues in IDRs predicted by ESpritz X-Ray (X-Ray), ESpritz Disprot (Disprot), and ESpritz NMR (NMR) in the "main" **(A)** and "additional" **(B)** datasets. Statistical analysis was performed using a hypergeometric test (*** P < 0.001).

### Supplementary Figure 11

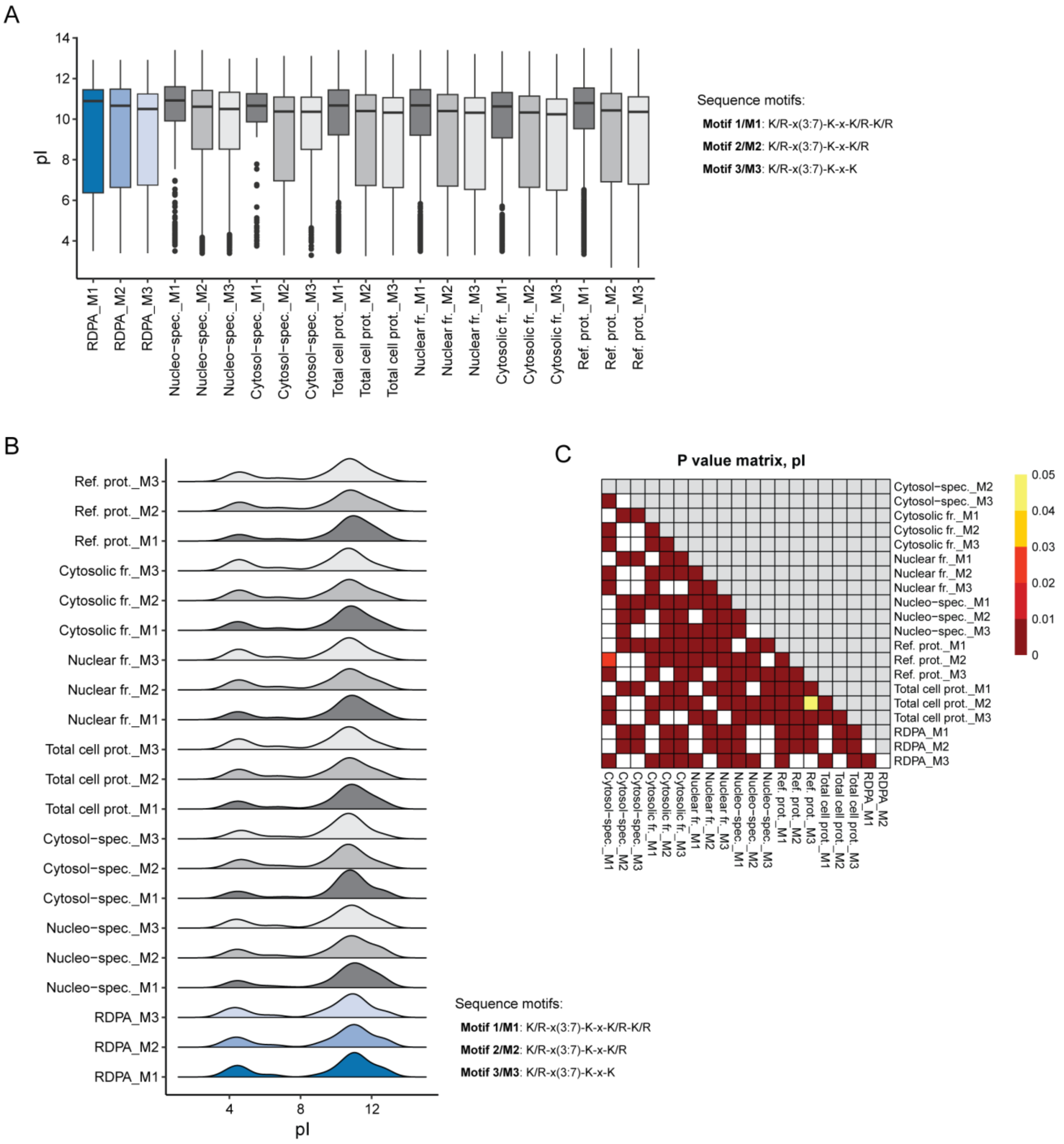


**Supplementary Figure 11: Additional bioinformatic analysis of RDPA proteome features (relevant to Figure 2H).** **A)** Boxplots show the distributions of pI values of IDRs between datasets and K/R motifs. **B)** Density plots of the IDR pI values highlight the presence of bimodal distributions. **C)** The P values of all pairwise comparisons between the datasets and motifs were estimated by a pairwise Wilcox test. Benjamini-Hochberg correction was applied to correct for multiple hypothesis testing. Ref. – reference, prot. – proteome, fr. – fraction, spec. – specific.

**Supplementary Figure 12**

A

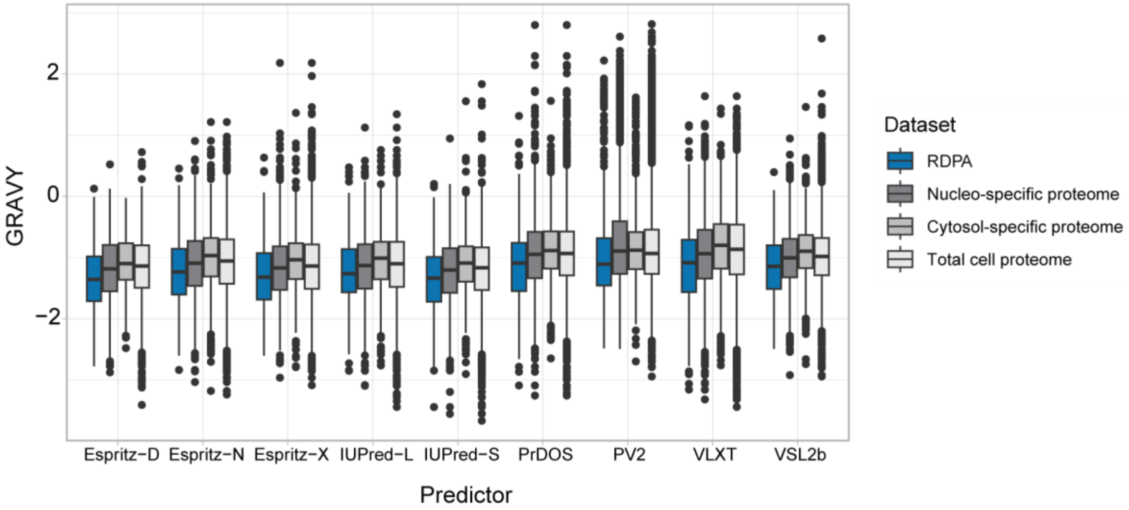


B

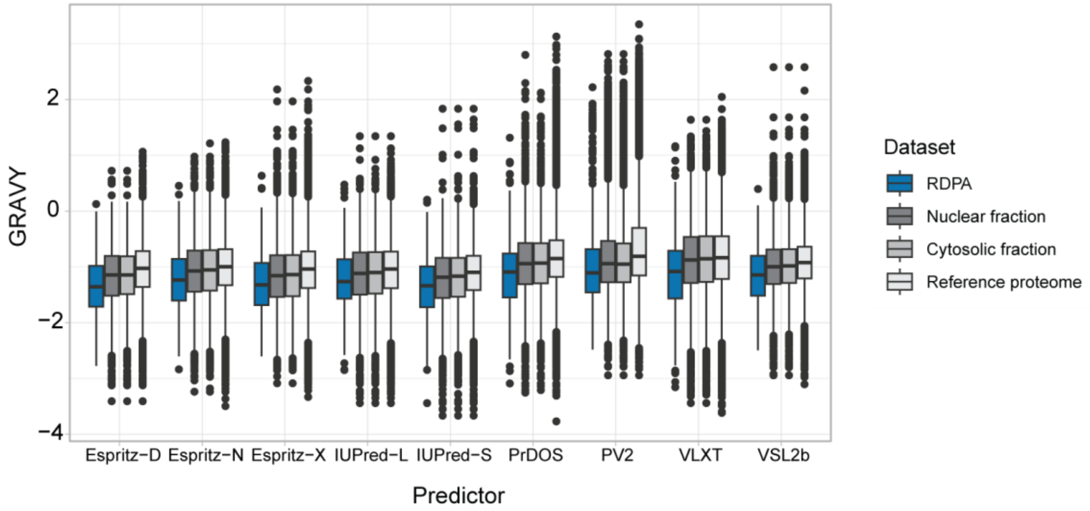


**Supplementary Figure 12: Additional bioinformatic analysis of RDPA proteome features (relevant to Figure 2I). A-B)** Distribution of the GRAVY scores of IDRs predicted by nine different IDR predictors (Database of Disordered Protein Predictions; only IDRs with minimal length of 20 amino acid residues were considered) in the "main" **(A)** and "additional" **(B)** datasets.

**Supplementary Figure 13**

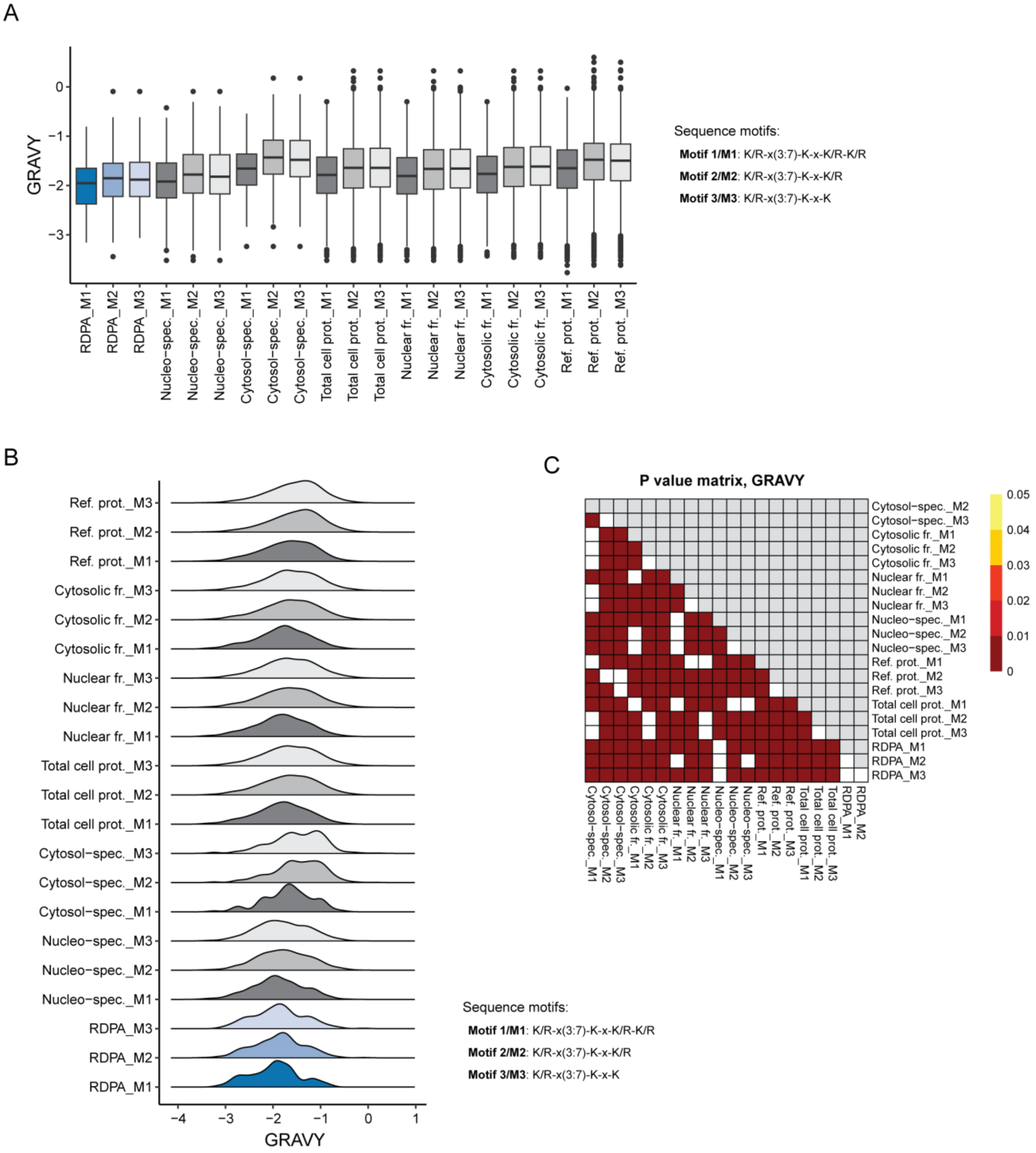


**Supplementary Figure 13: Additional bioinformatic analysis of RDPA proteome features (relevant to Figure 2I).** **A)** Boxplots show the distributions of the GRAVY score of IDRs between datasets and K/R motifs. **B)** Density plots of the IDR GRAVY scores. **C)** The P values of all pairwise comparisons between the datasets and motifs were estimated by a pairwise Wilcox test. Benjamini-Hochberg correction was applied to correct for multiple hypothesis testing. Ref. – reference, prot. – proteome, fr. – fraction, spec. – specific.

**Supplementary Figure 14**

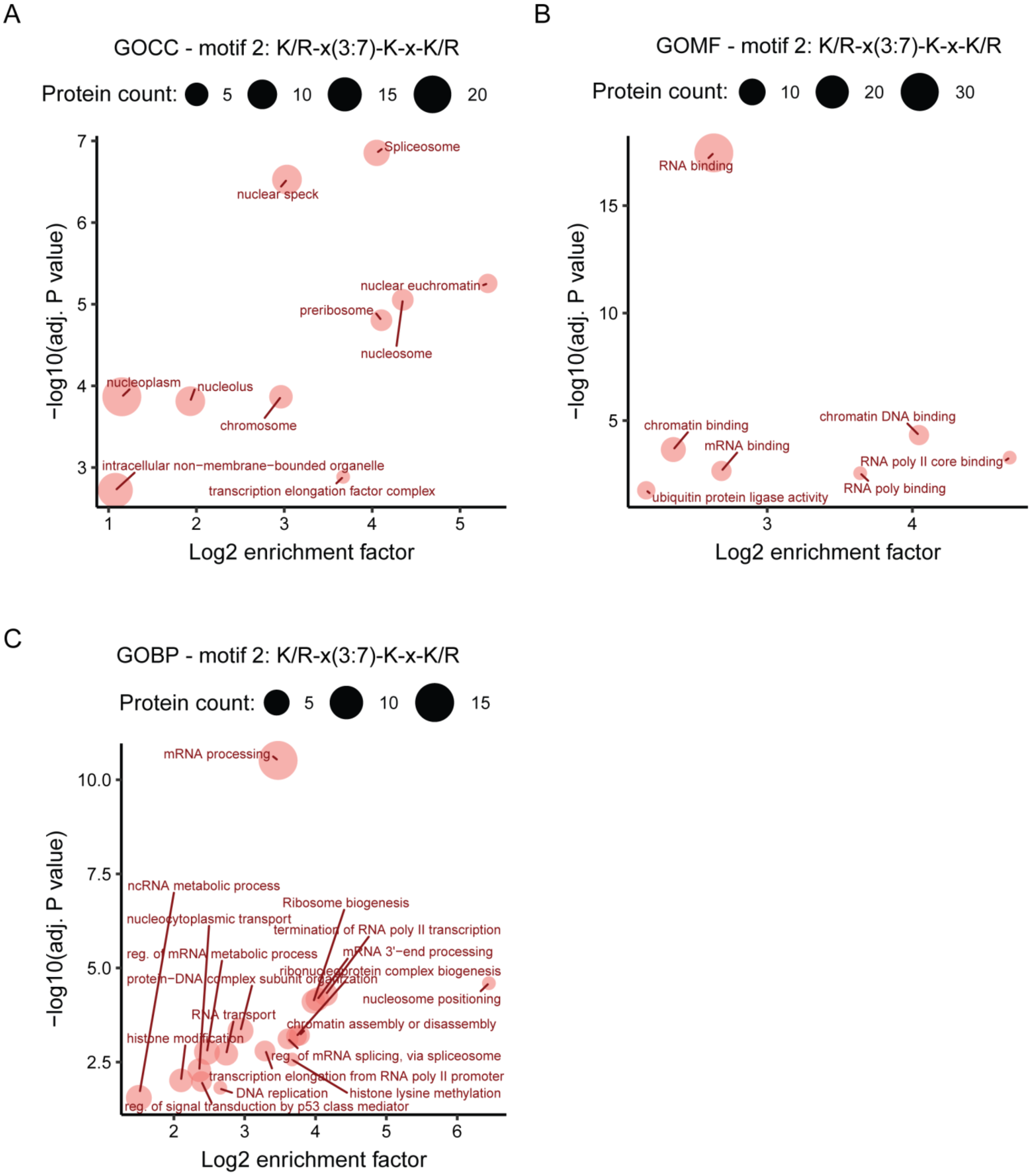


**Supplementary Figure 14: Functional analysis of the RDPA proteome (relevant to Figure 4B-D). A-C)** Gene ontology (GO) analysis of human proteins containing K/R-x(3,7)-K-x-K/R motif in IDRs using SLiMSearch tool based on **(A)**, cellular compartment (GOCC), **(B)** molecular function (GOMF), and **(C)** biological process (GOBP). The yaxis shows the –log10 adjusted p-value (Fisher's exact test) of proteins from a GO category, the x-axis shows the log2 enrichment factor. The size of the bubble corresponds to the number of proteins.

**Supplementary Figure 15**

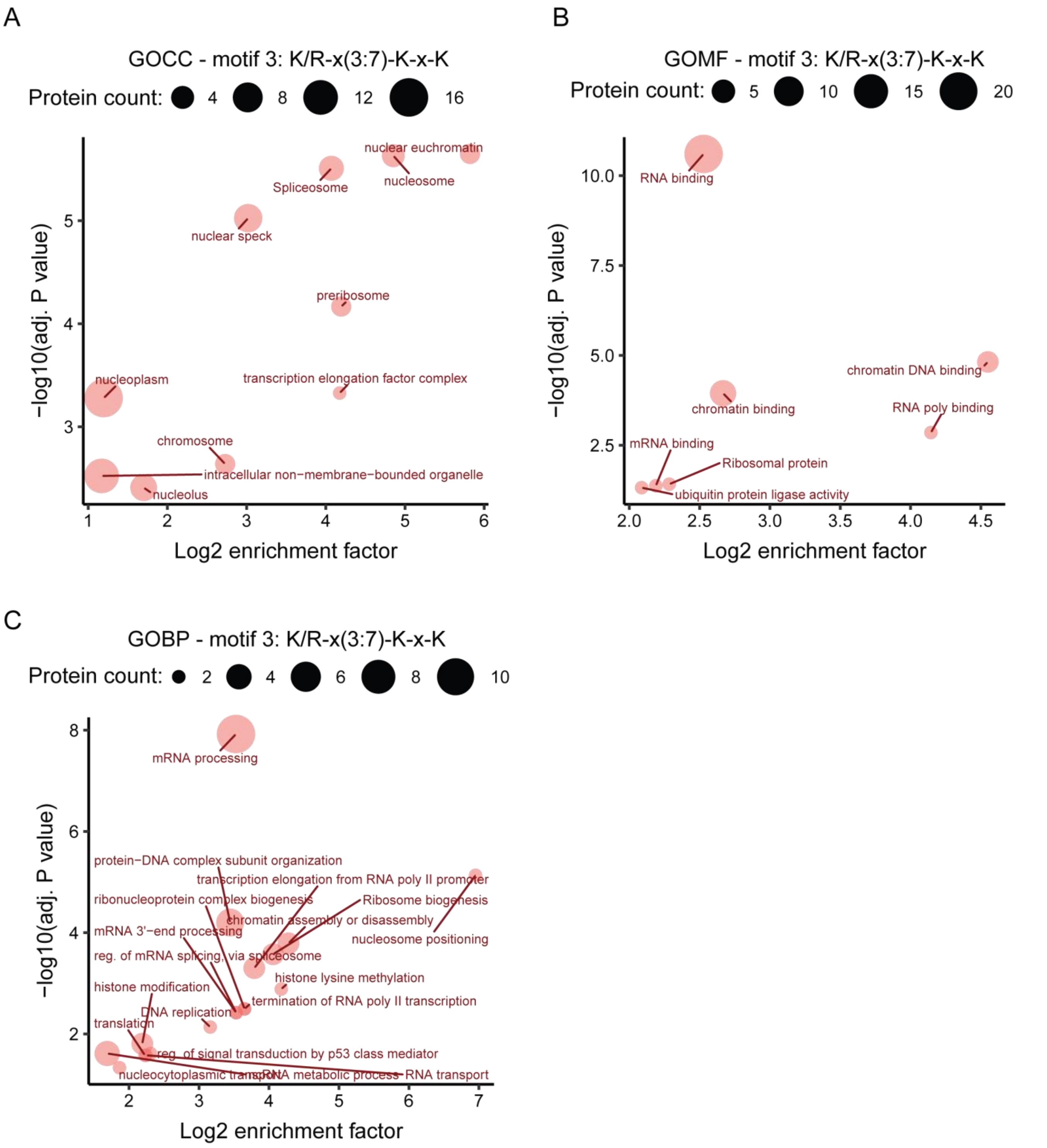


**Supplementary Figure 15: Functional analysis of the RDPA proteome (relevant to Figure 4B-D). A-C)** Gene ontology (GO) analysis of human proteins containing K/R-x(3,7)-K-x-K motif in IDRs using SLiMSearch tool based on **(A)**, cellular compartment (GOCC), , **(B)** molecular function (GOMF), and **(C)** biological process (GOBP). The yaxis shows the –log10 adjusted p-value (Fisher's exact test) of proteins from a GO category, the x-axis shows the log2 enrichment factor. The size of the bubble corresponds to the number of proteins.

**Supplementary Figure 16**

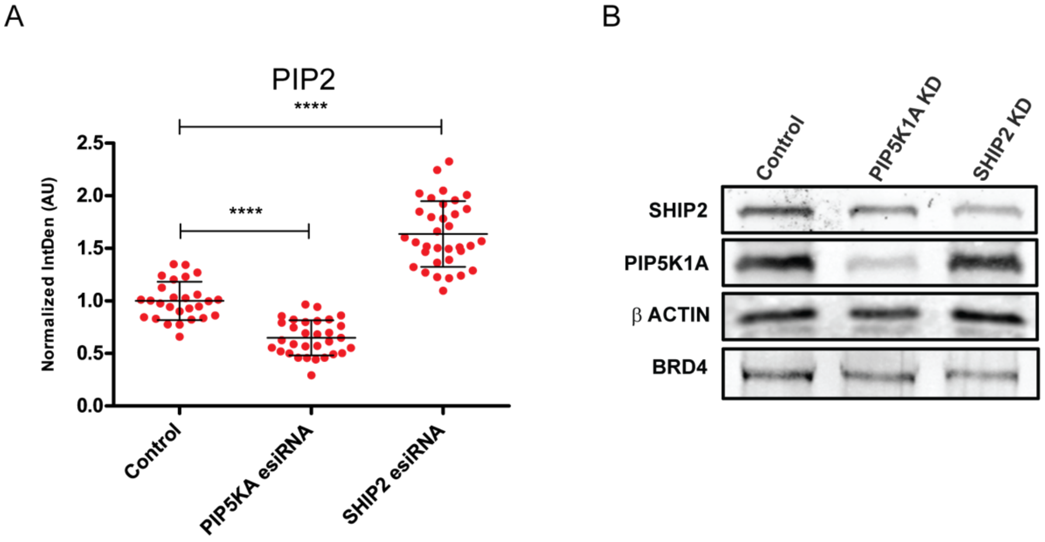


**Supplementary Figure 16: Manipulation of PIP2 level by PIP5KA and SHIP2 knock-down (relevant to Figure 5).** **(A)** Microscopy confirmation of the manipulation of PIP2 levels induced by depletion of PIP5KA and SHIP2 enzymes determined by microscopy. **(B)** WB analysis of the efficacy of PIP5KA and SHIP2 depletion and its effect on BRD4 protein levels.
